# LAGOS-US LANDSAT: Remotely sensed water quality estimates for U.S. lakes over 4 ha from 1984 to 2020

**DOI:** 10.1101/2024.05.10.593626

**Authors:** Patrick J. Hanly, Katherine E. Webster, Patricia A. Soranno

## Abstract

Broad-scale, long-term water quality (WQ) studies are critical for understanding increasing pressures on inland waters but remain rare due to cost. The LAGOS-US LANDSAT dataset provides 37-year remote sensing-derived WQ estimates for thousands of U.S. lakes ≥ 4 ha (1984–2020). WQ estimates use machine-learning models with Landsat imagery and ground-truthed Water Quality Portal data (LAGOS-US LIMNO). The dataset includes: (a) 45.9 million whole-lake water reflectance (six bands and 15 band ratios); (b) 740,627 matchups from 13,756 lakes with in situ data for six WQ variables: chlorophyll, Secchi depth, true color, dissolved organic carbon, total suspended solids, or turbidity; and (c) predictions for each WQ variable across lake-time combinations with quality imagery. Two random forest models were fit for each variable: Holdout-data (75/25 spatially representative train-test split) and Full-data (trained on all data). Variance explained for the Full-data predictions ranged from 20.7% for TSS to 63.7% for Secchi depth. Imagery underwent cloud and pixel quality control, and workflow components were validated guiding future research.

## Background & Summary

Contemporary studies of lake water quality (WQ) at regional, continental, and global scales are constrained by limited in situ monitoring data. This scarcity of comprehensive and consistent data hinders our ability to assess how WQ has responded to recent global changes, including growing pressures from land-use change and increasing climate variability. Additional approaches are needed to address these data gaps. We present LAGOS-US LANDSAT, a dataset that includes the WQ observations, and also the underlying data, full documentation, and model code to enable not only research using these data, but to enable the development of similar long term datasets in other locations using this method. Our dataset leverages the expanding Landsat image archive of remotely sensed images that can provide estimates of WQ in regions of the globe or time periods that have been poorly sampled. Our objective was to build a database of long-term observations of six water quality variables in tens of thousands of lakes ≥ 4 ha in the conterminous US from 1984 - 2020 using remotely sensed imagery from the Landsat SR data product. We used Landsat because it is the longest-running earth-observing satellite platform and has been used extensively in methods development for measuring inland WQ^1–8^.

Previous studies, particularly on US and Chinese lakes, have advanced methods for data integration and WQ modeling to leverage Landsat for broad-scale studies^1,7,11–13^ (**Table 1**). For example, AquaSat^7^ is an open-access dataset for the conterminous US containing match-ups between surface reflectances from Landsat SR and in situ WQ observations collected within one day. Predictive models for specific WQ components, such as Secchi depth^11^ and trophic categorization based on the Nutrient-Color Paradigm^13^, have also been created at the US national scale. These databases are supported by well-documented and fully reproducible methods that enable other researchers to both use their data and build new applications off them. Other broad-scale data products have more limited reusability, such as ground-truth Secchi depth predictions provided as 5-year averages for over 10,000 Chinese lakes across the Landsat platform period^14^. Additional global datasets that focus more on addressing scientific questions rather than producing reusable data products have been published for Secchi depth^15^ and for algal bloom metrics^15–18^. However, none of these studies provides long-term estimates of multiple WQ variables or for lakes as small as 4 ha.

**Table 1.** Synthesis of published studies that use Landsat for macroscale studies of lake water quality.

| Characteristics of macroscale lake studies using Landsat |  |  |  |  |  |  | OA data product includes |  |  |  |  |  |
| --- | --- | --- | --- | --- | --- | --- | --- | --- | --- | --- | --- | --- |
| Country or spatial extent | Name of resource | Temporal extent (maximum possible) | Min lake size | Number of lakes | Aquatic Variables of interest | Study type | Reflectances | Match-ups | Ground-truthed predictions | Calculated values from reflectances | Methods documentation | Reference |
| Global | NA | 1982-2018 | > 100 ha | 176,032 | Algal blooms (FAI) | Analysis : Calculations from reflectances |  |  |  | ✓ | No (Only methods section; no code) | Fang et al. 2022 |
| Global | Global Bloom Database | 1982-2019 | > 10 ha | 248,243 | Algal blooms (FAI) | Analysis : Calculations from reflectances |  |  |  | ✓ | No (Only methods section; no code) | Hou et al. 2022 |
| Global | NA | 2019 | > 1 ha | NA | Secchi | Model development |  |  |  |  | No (Only methods section; no code) | Song et al. 2022 |
| Global | NA | 1984-2022 | NA | > 300 | Secchi | Model development | ✓ |  | ✓ |  | Partial (Docum. & code available, but not reproducible) | Maciel et al. 2023 |
| China | Annual inland water clarity dataset of China | 1984-2018 (5-yr means) | > 1 ha | 10,814 | Secchi | Dataset: Ground-truthed predictions |  |  | ✓ |  | Partial | Tao et al. 2022 |
| China | CNLTSI | 1984-2023 | > 100 ha | 2,693 | Trophic State Index (TSI) | Dataset: |  |  | ✓ |  | Partial (Docum. & code available, but not reproducible) | Hu et al. 2024 |
| Continental US | AquaSat | 1984-2019 | NA | Sites, not lakes | TSS, DOC, CDOM, CHL, Secchi | Dataset: Reflectances (6 bands); Match-ups | ✓ | ✓ |  |  | ✓ | Ross et al. 2019 |
| Continental US | LimnoSat-US | 1984-2020 | > 10 ha | 56,792 | NA | Dataset: Reflectances (6 bands) | ✓ |  |  |  | ✓ | Topp et al. 2020 |
| Continental US | NA | 1984-2018; 5-yr means | > 10 ha | 14,000 | Secchi | Analysis : Ground-truthed predictions | ✓ |  | ✓ |  | ✓ | Topp et al. 2021 |
| Continental US | NA | 1984-2020 | > 10 ha | 26,000 | Spectral color | Analysis : Calculations | ✓ |  |  | ✓ | ✓ | Topp et al. 2021 |
| Continental US | LTS-US | 1984-2020 | > 10 ha | 55,662 | Trophic state | Dataset: Modeled predictions |  |  | ✓ |  | ✓ | Meyer et al. 2023 |
| Continental US | LAGOS-US LANDSAT | 1984-2020 | > 4 ha | 106,000 | Secchi, CHL, DOC, True color, TSS, Turbidity | Dataset: Reflectances (6 bands); matchups; ground-truthed predictions | ✓ | ✓ | ✓ |  | ✓ | This study |

Our study and data release^19^ builds on this prior work to advance the use of Landsat SR for broad-scale studies of inland waters. Although using Landsat and ground-truthed WQ observations to predict WQ in unsampled lakes has a long history, many workflow components remain untested at the scale of thousands of lakes. We present results from these tests to guide future applications. Our advancements include: (1) testing computationally expensive components of the workflow to determine when and where they are needed to improve predictions; (2) creating a single data product with six WQ variables in addition to the reflectances, match-ups with in situ data, ground-truthed predictions, and data quality flags; and, (3) creating and testing the validity of using Landsat SR for a broader range of WQ variables. Our final data product meets the FAIR data principles of findability, accessibility, interoperability, and reusability (20) and we include validation of predictions, methods, and recommendations for specific processing steps.

Our dataset provides ground-truthed estimates for six WQ variables from 1984 to 2020, derived from over 45 million images from Landsat 5, 7, and 8 for 136,977 lakes, with a median of over 121,000 lakes sampled annually. In addition to the WQ estimates, we include: five data flags for users; reflectance values on all individual bands and 2-band ratios for all lake-date combinations; Landsat platform identifiers and quality metrics; and our matchup dataset that includes matchups within 0 to 7 days in either direction. The dataset is fully interoperable with the LAGOS-US research platform, which provides millions of data points on hundreds of lake features relevant to WQ including delineated watersheds, regional characteristics, land use, lake depth, and other variables^21–23^.

The six WQ variables are chlorophyll a (CHL), Secchi depth, true color, dissolved organic carbon (DOC), total suspended solids (TSS), and turbidity. Chlorophyll a (CHL; μg/L) estimates suspended phytoplankton biomass in the pelagic zone, corrected for pheophytin, serving as a key measure of lake productivity. Secchi depth (meters) measures water clarity using a disk lowered until no longer visible.

True water color (true color) measures the colored fractions of dissolved organic material using filtered water samples, reported in platinum cobalt units (PCU). Dissolved organic carbon (DOC; mg/L) measured from filtered water, includes both colored and non-colored fractions of organic carbon. Total suspended solids (TSS; mg/L) include silt, clay, sand, algae, and plant material suspended in water.

Turbidity, another measure of water clarity, measures light scattering due to suspended particles and is measured in Nephelometric Turbidity Units (NTU). Together, these variables provide a comprehensive snapshot of WQ status and biological functioning in inland waters, reflecting eutrophication pressures, land-water interactions, climate change, hydrologic forcing, and food web dynamics. While CHL, true color, DOC, TSS, and turbidity represent specific components of dissolved or suspended particulate matter, Secchi depth provides an integrated measure of all components. Full definitions, source information, and method descriptions for these six variables are described in the documentation of methods for the LAGOS-US LIMNO data product we used for our in situ WQ data^10^.

## Methods

We developed a reproducible workflow for generating long-term, remotely sensed water quality (WQ) predictions for 136,977 lakes ≥ 4 hectares across the conterminous United States (1984–2020). Following established approaches, we used the Landsat Surface Reflectance (SR) data product to develop ground-truthed WQ predictions over time. We used Google Earth Engine (GEE) to extract reflectances for over 100,000 lakes across the 37-year period. Because many methodological components for inland water remote sensing remain untested, we evaluated several key workflow components to guide future researchers using Landsat SR data in GEE for large-scale WQ predictions. Our overall approach, illustrated in **Figure 1** for an individual WQ variable, was repeated for all six variables. Detailed methods for each step follow.

**Figure 1.**
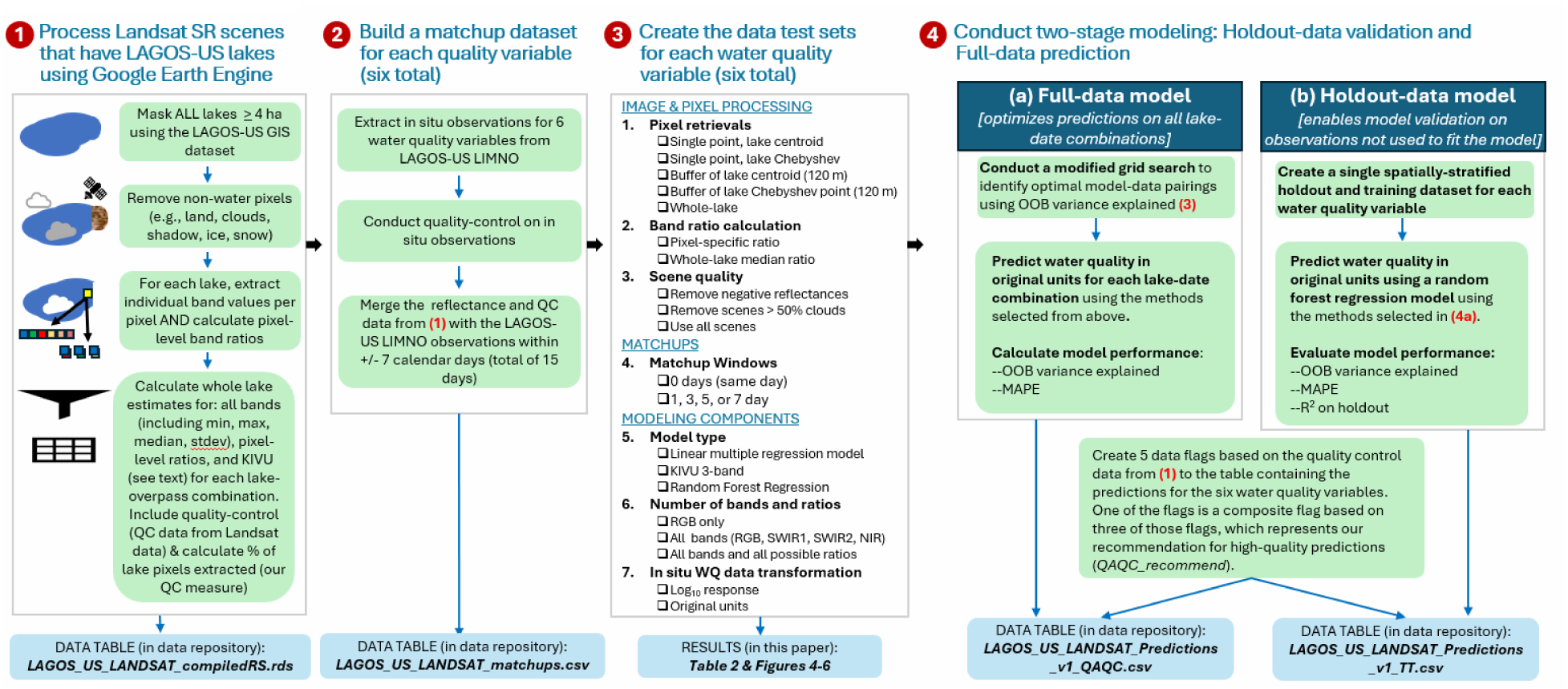
Overview of the four major steps used to create LAGOS-US LANDSAT. See text for full description of all steps. The steps are described for an individual water quality variable (e.g., CHL). Because the same dataset in step 1 is used for all models, steps 2-4 are repeated for each water quality variable.

### (1) Process Landsat SR scenes

We used the Tier 1, Level-2 Landsat Surface Reflectance product (Collection 1), which employs the Landsat Ecosystem Disturbance Adaptive Processing System (LEDAPS)^24^ to atmospherically correct scenes from Landsat 5 (TM) and 7 (ETM+) and the Land Surface Reflectance Code (LaSRC)^25^ to atmospherically correct scenes from Landsat 8 (OLI). We extracted surface reflectance values for all 137,465 LAGOS-US lakes ≥ 4 hectares in surface area^21^, from 1984-2020 (1985-2020 complete years) using GEE (**Figure 1**, step 1). We selected 4 ha as our minimum lake size because any smaller area results in large numbers of lakes with pixels that have mixed water and land components, making it challenging to capture true water-only pixels. Because lakes were run as separate instances and the default average concurrent tasks allowed on GEE at the time of execution was two, the total runtime for all lakes took over a year to complete and increases substantially with lake size in our method where all pixels in a polygon are extracted.

For each lake and for each sensor overpass and band, we extracted the median, minimum, and standard deviation of individual pixel reflectance values, and band ratios calculated at the per-pixel level before a lake-wide median was extracted. The potential area for extraction was fixed to the static polygons from LAGOS-US LOCUS for the full time series. We removed pixels coded as clouds, cloud shadows, snow/ice, or fill data, retaining only clear-sky pixels over water using the CFMASK-derived pixel quality attributes^26^ in the Landsat Pixel Quality Assessment Band. To enable historical time series analyses across the full study period we harmonized bands across the TM, ETM+, and OLI sensors, for which we used complete handoff following the launch of a new sensor, using published surface reflectance transformations^6^ prior to reflectance extraction.

As part of our data product, we created the data table, *LAGOS_US_LANDSAT_compiledRS*, which contains output from the steps described in **Figure 1a** to aid in later steps and to make data available to other users for their modeling purposes in the future. Each row in the data table represents a unique lake-scene combination and includes: median lake-wide values for the 6 bands available for all sensors, pixel-specific ratios for all possible band combinations (n=15), the percent cloud cover of each scene, the total number of pixels extracted, and a 3BDA-like “KIVU” algorithm applied to the individual pixels^27^. We also created data flags for: (1) any pixel or any band with negative reflectances; (2) any lake median reflectance that is negative; (3) the percent pixels retrieved for a lake; and (4) duplicate scenes on the same day for a lake (rare, but present due to overlapping overpasses). Our final data product represents 136,977 lakes (99.6% of LAGOS-US lakes ≥ 4 hectares) that had at least one set of Landsat reflectances and provides 45,867,023 reflectance sets of lake and overpass combinations through time.

We retrieved an average of 48.0% of all pixels within a lake polygon, with 37,529,125 lake-scene combinations meeting or exceeding our 10 percent pixel threshold (81.8%). Finally, 4,877,886 lake reflectance sets had shared calendar days for a lake due to scene overlap (1.1%).

### (2) Build a matchup dataset for six water quality variables

Our matchup dataset **(Figure 1, step 2)** is based on the LAGOS-US LIMNO data product^10^ that contains in situ data extracted primarily from the Water Quality Portal and three sampling years from the USEPA National Lakes Assessment. We conducted quality control analyses on each of the six WQ variable matchup sets and made the following data decision. First, for CHL, we removed negative values and extremely large values > 800 ug/L that are rarely measured in pelagic zones of lakes. Second, if a lake had multiple measurements for a given WQ variable on a calendar day, the median value was calculated prior to matching with remote sensing data. Third, because the few observations for true color > 100 PCU appeared to skew the distribution and were from a small number of primary data sources, we removed them from the training set, recognizing that our resulting models would likely not be able to capture the most extreme high range of true color. Nevertheless, because lakes are considered tannic if they have values > 20 PCU^28^ and highly colored lakes were rare, we made the decision to apply a data limit in order to produce more accurate training models. The ranges of data for all other variables were less skewed and were not trimmed.

After merging the remote sensing dataset with the LIMNO dataset on a calendar date basis (with a maximum matchup window of +/- 7 days), we had 723,206 matchups of lake-date combinations with at least one observation for any of the six WQ variables across 13,756 LAGOS-US lakes. The number of matchups (including multiple values by date per lake) by WQ variable were: CHL = 238,248; Secchi depth = 666,060, DOC = 8,269; true color = 37,839; TSS = 52,677; turbidity = 68,573. Matchup lakes were widely distributed across the conterminous US with high densities of observations related to regional lake density and monitoring program specifications (**Figure 2**). Although substantial matchup data exist across the study period, changing monitoring intensity and resulted in an uneven temporal distribution, with matchup numbers peaking around 2008 (**Figure 3**). The complete set of matchup data that are within +/- 7 days is provided in the data table *LAGOS_US_LANDSAT_matchups*.

**Figure 2.**
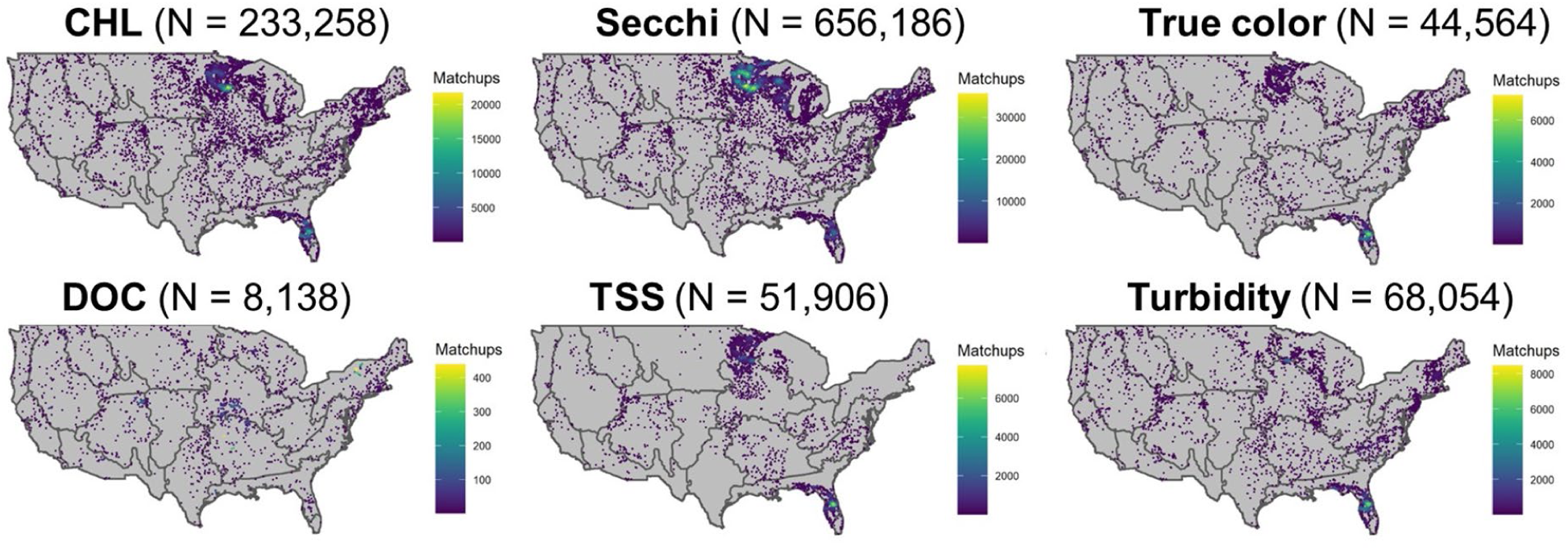
Maps of the location of lakes in the LAGOS-US LANDSAT matchup dataset by water quality variable. For display purposes, we show the matchup data that are within ±7 calendar days of the LAGOS-US LIMNO in situ observations for each of the six water quality variables which represents the maximum possible matchup dataset for each variable and the maximum sample sizes and what was tested in our modeling workflow. See results in Table 2 for the final matchup windows used for each variable as well as the final sample sizes used. The map outlines are NEON region borders.

**Figure 3.**
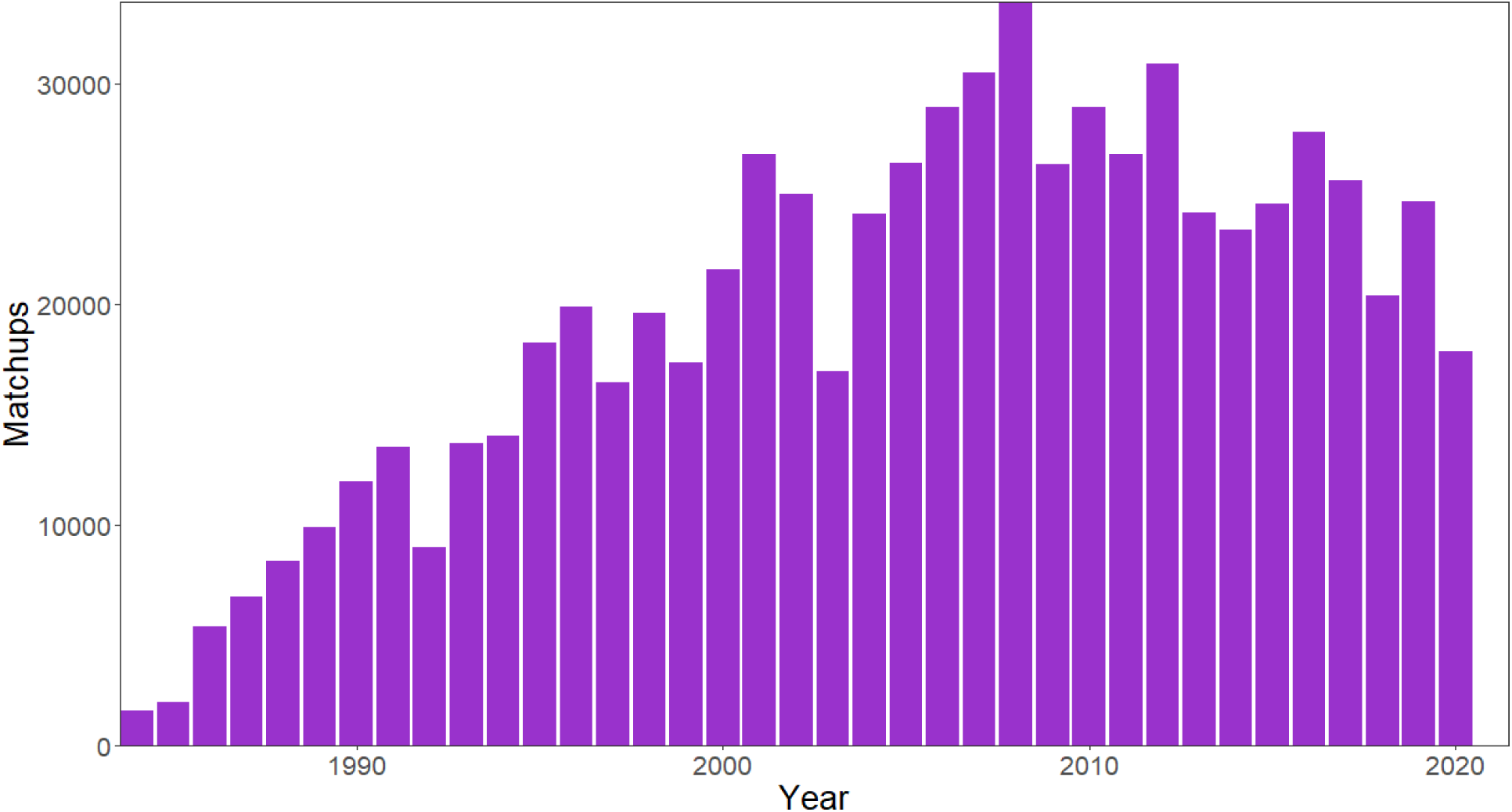
Histogram of the number of matchups by year for the matchup dataset. The matchup window includes all dates within ±7 calendar days of the LAGOS-US LIMNO in situ observations for all water quality variables as a sum across all variables within each year.

**Table 2.** LAGOS-US LANDSAT workflow decisions and fit statistics for the Full-data model for each water quality predictive model.

| *Workflow components tested and selected* |  |  |  |  |  |  |  |  |  | *Final model fit statistics* |  |  |  |  |
| --- | --- | --- | --- | --- | --- | --- | --- | --- | --- | --- | --- | --- | --- | --- |
| Image and pixel processing by lake |  |  |  |  | Modeling steps |  |  | Matchup training data |  |  |  |  |  |  |
| Variable | Pixel retrievals | Band ratio calculation | Scene quality: Negative reflectances | Scene quality: Scene clouds > 50% | In situ data transformed | # of bands and ratios | Training model type | Matchup window (± days) | Final matchups (N) | Variance explained (%) | RMSE | MAPE (%) | MAPE (%) Improvement of $\log_{10}$ model | |
| CHL | Whole-lake | Pixel-specific | Removed | Removed | Log<sub>10</sub> | All bands & ratios | Random forest | 1 | 43,755 | 44.1 | 0.43 | 23.3 | (+9.8) |  |
| Secchi | Whole-lake | Pixel-specific | Removed | Removed | none | All bands & ratios | Random forest | 7 | 586,368 | 63.7 | 1.23 | 22.8 | -- |  |
| True color | Whole-lake | Pixel-specific | Removed | Removed | Log<sub>10</sub> | All bands & ratios | Random forest | 7 | 33,194 | 48.6 | 0.26 | 7.5 | (+12.7) |  |
| DOC | Whole-lake | Pixel-specific | Retained | Retained | Log<sub>10</sub> | All bands & ratios | Random forest | 7 | 7,466 | 42.0 | 0.24 | 10.2 | (+6.1) |  |
| TSS | Whole-lake | Pixel-specific | Removed | Removed | Log<sub>10</sub> | All bands & ratios | Random forest | 1 | 10,845 | 20.7 | 0.62 | 27.0 | (+16.3) |  |
| Turbidity | Whole-lake | Pixel-specific | Removed | Removed | Log<sub>10</sub> | All bands & ratios | Random forest | 1 | 58,999 | 52.1 | 0.37 | 8.8 | (+20.9) |  |

### (3) Create the data test sets for each water quality variable

We tested major methodological components related to image and pixel processing, modeling, and matchup windows (**Figure 1, step 3**). Using the matchup dataset, we conducted exploratory assessments of key workflow components for training models for all six water quality variables. Because methodological decisions were interrelated, analyses were not performed in linear order. While most tests included all water quality variables, computational constraints in GEE limited some tests to one to three variables. For example, using a randomly selected lake matchup dataset of 500 lakes with in situ DOC data, we ran a set of 40 training models that varied the image and pixel processing steps, transformations, and matchup windows. In addition to identifying the best-performing model type (**Figure 4**), we tested the influence on prediction quality for the following model inputs (**See Figure 1, step 3**): 5 types of pixel retrievals; 2 types of band ratio calculations; 2 types of scene quality indicators; and inclusion of all bands or all bands plus all ratios. Because many components that we tested have not been analyzed in any published studies to date, the outcomes of these tests are informative not only for our final model selection but also for future studies using the Landsat SR platform for broadscale water quality studies.

**Figure 4.**
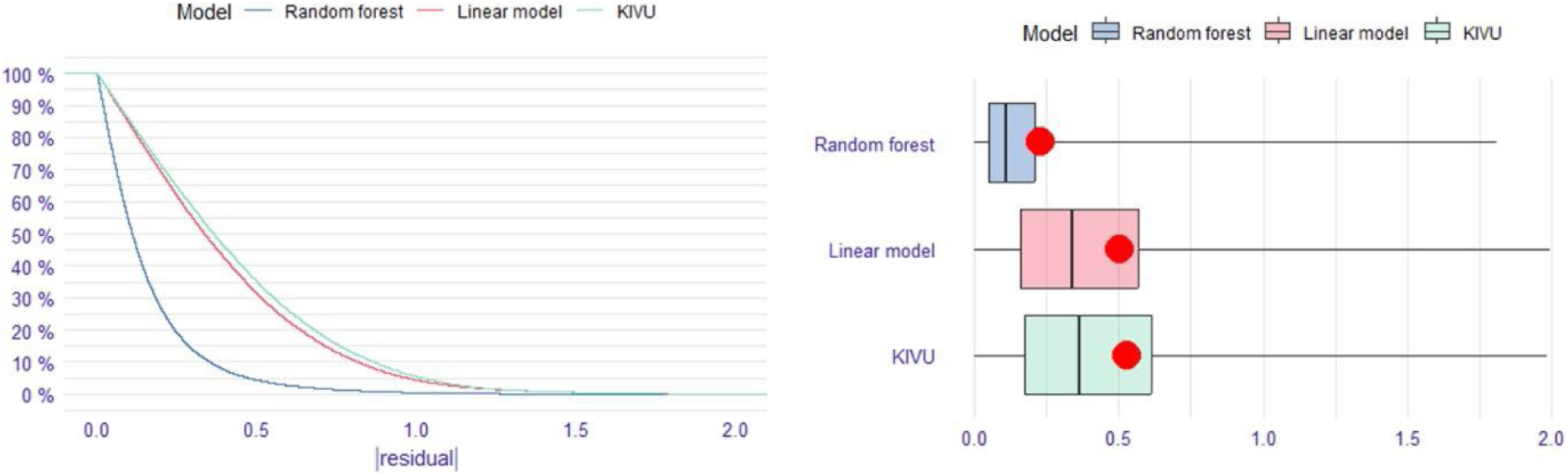
Results from the three model approaches tested for the CHL model. The left panel shows the reverse cumulative distribution of the absolute value of residuals of the three models using log10-transformed in situ CHL data and using matchups within ±1 day. See text for model details. ‘Random forest’ is a random forest regression model that includes all bands and band ratios; ‘Linear model’ is a linear multiple regression model with all bands and band ratios; and ‘KIVU’ is a 3-band algorithm calculated at the pixel level for each lake. The right panel shows boxplots of these same absolute values of residuals with the red dot indicating the RMSE: random forest = 0.23, linear model = 0.51, ‘KIVU’ = 0.54.

**Table 2** summarizes seven main outcomes from our workflow tests for building large-scale remote-sensing models using the Landsat platform. All tests were conducted for all water quality variables and lakes, except the first two steps, which were either computationally prohibitive or unnecessary for the full dataset.

#### Image and pixel processing

1. **Pixel-retrieval calculated at the whole-lake scale performed best**. The lake center with a buffer performed close to the whole-lake estimates while the lake center calculated using the Chebyshev point was consistently better than the lake centroid (**Figure 5a**). This test was extremely resource-intensive and so we only tested it for a subset of lakes and for a subset of water quality variables (CHL, Secchi depth, DOC).

**Figure 5.**
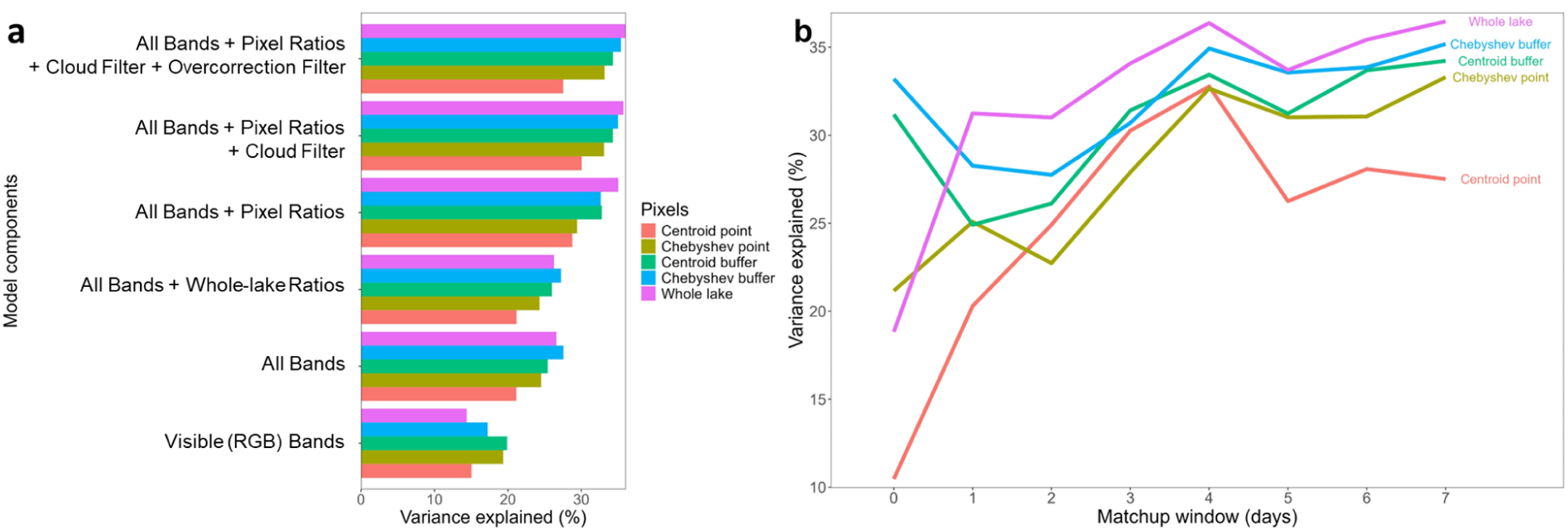
Results from testing different components of the workflow related to image and pixel processing for a subset of the DOC dataset. (a) A plot of variance explained for each training model run conducted on log10-transformed in situ data that included the indicated components. Results are plotted for the 5 different ways that reflectance is summarized by lake (see text). ‘Whole-lake Ratios’ are ratios calculated on median band data, whereas ‘Pixel Ratios’ are those ratios first calculated at the pixel-level and then the median value is calculated; ‘All Bands’ includes all bands; ‘Overcorrection Filter’ is the flag for when there is negative median reflectance. (b) A plot of % OOB variance explained by matchup window and the 5 different ways that reflectance is summarized by lake. Models are based on a 500-lake subset that contained DOC matchups, with some lakes being sampled more than once (N = 1,348).

2. **Calculating band ratios at the pixel level was always better,** i.e., with lower prediction error, than first calculating lake-wide or buffer-wide band averages and then calculating ratios from those (Figure 5a). Because this step must also be conducted within GEE, it was critical for consideration early in the workflow.

3. **Removal of negative reflectances and scenes > 50% clouds was best (except for DOC).** The results for DOC were likely due to the lower sample size of the matchup data. DOC (**Table 2**; **Figure 5a**). However, the differences in this test were not large, so when sample sizes are low, retaining some scenes with poorer quality may be better because sample size is increased. In general, these poor-quality images were relatively rare, so the gain in predictive performance was not large across many lakes.

#### Matchups

4. **The optimal matchup window ranged from 1 to 7 days across the WQ variables and should always be tested.** We found the optimal matchup window differed by WQ variable and was likely due to the inherent daily variability in the WQ variable (**Table 2**; **Figure 5b)**. For example, true color is a more temporally stable variable for many lakes than CHL, TSS, and turbidity which can all respond readily to large precipitation or wind events day to day. Consequently, the optimal matchup window for the latter 3 variables was 1 day but was 7 days for Secchi depth, true color, and DOC. Nevertheless, we encourage users to test this important part of the workflow in future applications, and the results may be specific to the type of lakes being modeled.

#### Modeling components

5. **The machine learning model performed best**. We tested three types of models that have been used by other studies using the CHL dataset: linear models with all bands plus band ratios; the three-band KIVU retrieval algorithm^29^ that was previously found to be relatively effective for inland water chlorophyll retrieval^27^; and random forest regression models using all bands plus band ratios. We found that the random forest regression model, applied using the ‘randomForest’ package in R^30^ far outperformed the other two models with over two times reduction in error compared to the other two models (**Figure 4**). We found similar gains in performance for Secchi depth data (data not shown). Based on this test that was conducted only for CHL and that is backed by prior studies demonstrating the superior performance of machine learning models^11^, all of the remaining tests were performed only using the machine learning model.

6. **Using all bands and all band ratios outperformed RGB-only models**. Although we present initial results only for DOC (**Figure 5a**), we retained the testing of these model components throughout the process. The variable importance plots for the final models for all WQ variables demonstrate that although the non-RGB bands were never the top predictor, ratios containing one of these bands were often the second or third most important variable (see **Figure 6**).

**Figure 6.**
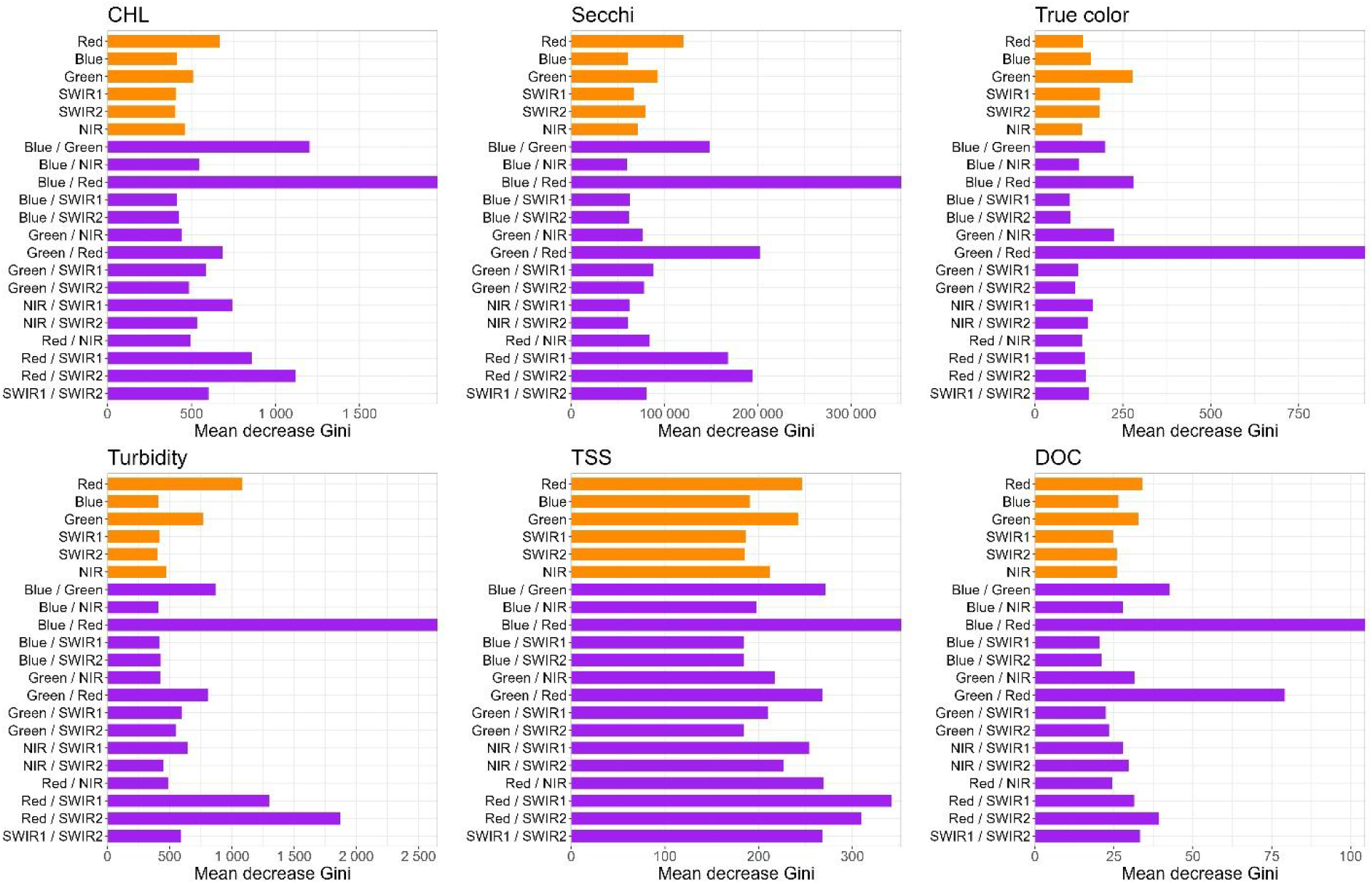
The full-data model variable importance (mean decrease Gini) for all predictors in the random forest regression models for each water quality variable. Orange bars are individual bands, and purple bars are ratios between two bands.

7. **Log-transformed in situ data resulted in better-performing models** than their non-transformed versions (**Table 2**, final column). Because the distributions of most of our WQ variables were left-skewed and more data-rich in the lower ranges, the models performed better when the transformations resulted in greater granularity at the lower ends of the data distributions.

### (4) Conduct two-stage modeling: Holdout-data validation and Full-data prediction

We implemented a two-stage modeling approach to balance the needs of different users while maximizing predictive accuracy. This dual approach provides both traditional error metrics through a spatially-stratified holdout validation procedure and optimal predictions using all available data. While we recommend the Full-data model predictions for most applications due to their superior coverage of spatial and temporal variability, we recognize that some users require traditional train-test validation metrics for comparison with other studies or regulatory compliance.

For our first modeling approach, we created a Holdout-data model test set based on a 75% training and 25% test split to provide external validation metrics. We used the Generalized Random Tessellation Stratified (GRTS) sampling method, implemented in the ‘spsurvey’ R package^31^ to create spatially representative splits. For our second modeling approach, we constructed a Full-data model leveraging the entire in situ dataset to maximize predictive accuracy across the full range of spatial and temporal variability in lake water quality. Because random forest models internally use out-of-bag (OOB) samples to assess prediction error, we used OOB variance explained as our primary performance metric. Prior studies have shown that OOB estimates provide a reliable alternative to test sets when sufficient trees are grown^32–33^. This method avoids sensitivity to a single train-test split, a known issue in ecological modeling^34^. We recommend the Full-data model and its OOB variance-explained for most ecological applications. The Holdout-data model is provided for users requiring traditional train-test error metrics.

Both modeling approaches followed identical workflow steps for each WQ variable. Our approach used a modified grid search to explore combinations of data filtering and workflow components, aiming to identify optimal model–data pairings for WQ prediction. We fit random forest regression models to each WQ Full-data dataset. The components varied included: matchup window (0, 1, 3, 5, or 7 days); modeling inputs (RGB, all bands, all bands plus whole-lake ratios, all bands plus pixel ratios); and data transformation (original units or log₁₀-transformed). After selecting the best-performing model from these 40 random forest models based on out-of-bag (OOB) variance explained we sequentially applied two additional filters: first, removing high cloud-cover scenes from the training data; and second, excluding matchups with any negative reflectance values in any band. In total, 42 models were fit separately for each water quality variable. The best-performing model for each variable was then selected to generate the final predictions. We then applied the same data filtering and modeling components within the Holdout-data model to better characterize the performance of the Full-data model for each WQ variable.

**Table 2** details the final model specifications and workflow steps with data decisions specified in each row. Final models were based on in situ training data that were log10-transformed (except for Secchi depth). Variance explained of predicted versus in situ data, a valuable model diagnostic, ranged from a low of 20.7% for TSS to a high of 63.7% for Secchi depth, ranging between 42-52% for the remaining WQ variables. Thus, except for TSS, the variance explained is quite high and certainly suitable for broad-scale water quality studies. All random forest models included all bands and ratios. We found that the top predictors were always a 2-band ratio based on variable importance factors calculated for each WQ model (**Figure 6)** like other studies demonstrating the importance of the blue/red ratio and blue/green ratios for Secchi depth and CHL. However, fewer studies have also included the SWIR bands, which were also quite important. Interestingly, the top predictor for true color was the Green/Red ratio, which was also the second highest ratio for closely related DOC, although, for DOC, the top predictor was Blue/Red, which was also top for Secchi depth and CHL. These results make ecological sense since DOC is related to both true color (which is the colored fraction of DOC) as well as CHL which is known to track with the non-colored fraction of DOC. Turbidity and TSS have similar dominant predictors, although the contributions of the other wavelengths and ratios differ suggesting they reflect distinct WQ features.

Our data product includes five quality-control flags, enabling users to filter data according to their study requirements:

1. Negative reflectance (any band) - flags lake-date combination where any band has negative value for at least one pixel
2. Negative median lake reflectance (any band) - flags lake-date combination where any whole-lake retrieval has a negative median value
3. Percent of lake pixels with quality reflectances - flags scenes with <10% of maximum lake pixels containing quality reflectances (e.g., due to cloud cover)
4. Duplicate predictions - identifies calendar dates with two satellite overpasses resulting in multiple predictions
5. QAQC_recommend - integrates the previous flags (2-4), removing data with negative median reflectance values and selecting the duplicate date prediction with the greatest number of pixels, to identify highest quality predictions that users can easily query to easily meet these criteria

## Data Records

The LAGOS-US LANDSAT data product^19^ consists of the following four data tables that include all components of our workflow described in **Figure 1**.

### LAGOS_US_LANDSAT_data_description.csv

This table is the combined data dictionary for the three tables below that describes and defines variables included in each of the four data tables. For each variable, we include the variable name, description, data type, and column index.

### LAGOS_US_LANDSAT_compiledRS.rds

This data table, in the form of an R Data Serialization (RDS) file, provides reflectance extraction output generated from Google Earth Engine that was extracted from a single lake at a specific time. Each row is a single LAGOS lake and Landsat scene combination. This data table contains 57 variables related to whole-lake summaries of reflectance values calculated for lake water pixels on all individual bands (min, max, median, standard deviation) and all 2-band ratios in addition to the KIVU calculation. Ratios are first calculated at the pixel level (referred to as pixel-wise in the data dictionary), and then the median is calculated across all lake-water pixels. We also include scene identifier information from the Landsat platform (satellite name, sensing time, scene ID, etc), quality estimates obtained from the Landsat platform (e.g., % cloud cover of the scene, image quality indicators), earth-sun distance, solar angles, etc.

### LAGOS_US_LANDSAT_matchups.csv

This data table includes all possible matchups within 0 to +/- 7 days between Landsat overpasses and in situ data obtained from the LAGOS-US LIMNO^10^ data module for the six WQ variables (CHL, Secchi depth, True color, DOC, TSS, Turbidity). For reference, we include the WQ variables from individual dates and the associated reflectance values and scene identifier information from the above dataset. If any date had more than one WQ estimate, we took the median value, however, more than one estimate was extremely rare, and the vast majority of data were from a single sampling event per day. We also include the matchup window value for each WQ in situ value so that future users can select the matchup window. Note that not all WQ variables are sampled on the same dates, so the sample sizes vary by WQ variable.

### LAGOS_US_LANDSAT_Predictions_v1_QAQC.csv

The predicted WQ data from the **Full-data model** that uses all available data for predictions for the selected matchup window. As described above in step 4 of our workflow, we include five data flags including the field QAQC_recommend in which we apply our criteria for quality predictions that users can select for a highest-quality version of the dataset. This dataset includes predictions using all data within the model-selected matchup window to minimize prediction errors.

### LAGOS_US_LANDSAT_TT.zip

The predicted WQ data from a **Holdout-data model** per WQ variable that is based on 75% train and 25% test datasets that is spatially balanced using the Generalized Random Tessellation Stratified (GRTS) algorithm. This dataset has predictions alongside measurements of prediction error from the training datasets using OOB variance explained and MAPE on the test datasets for each WQ variable. These datasets are useful for creating comparisons with other machine learning algorithms and for the generation of test data error estimates that are unseen by any tree in the modeling process.

### Technical Validation

The validity of using Landsat 5, 7, and/or 8 sensors for inferring water quality in lakes has been supported by decades of studies conducted on individual and combination of sensors^3,11,35–39^. In addition, the Landsat Collection 1 SR product has been noted for its suitability for machine learning estimation of CHL across the US^40^. However, challenges remain that careful methods development can address. For example, the spectral signals of WQ variables are not always independent; for example, colored dissolved organic matter and chlorophyll have overlapping absorption spectra^41^. Further, Landsat imagery may have lower accuracy for estimating WQ values in oligotrophic relative to eutrophic waters^42^. Therefore, we used a series of analyses to characterize the level of uncertainty in the LAGOS-US Landsat data and identify potential biases. In addition to quantifying numerous fit statistics and conducting validation that supports the use of the six models that we developed (**Table 2**), we also investigated the ecological interpretability of the predicted water quality estimates such as by comparing in situ and predicted individual lake time series.

### Model prediction error

In addition to examining the overall model fits that quantify model error across the whole dataset, we examine quantiles of predictions to examine whether errors differ across the range of the matchup dataset for both the Holdout-data (**Figure 7**) and Full-data (**Figure 8**) models. We found that in most comparisons, 90% of the data were close to the 1:1 line and that differences between predicted and in situ data are not that far off ecologically, especially when one considers predictions of water quality were made in over 100,000 lakes across 34 years. The model with the poorest performance was TSS (**Table 2**), which had the largest bias in predictions, with the median TSS being underpredicted relative to in situ data. Plots of error by binned values of predicted values also show that TSS has higher errors, particularly for high TSS lakes. We also found error by binned values for DOC to also be particularly high for the highest DOC values (> 16 mg/L). A study of global lake DOC estimates predicted the average DOC was 3.9 mg/L, with a maximum value of 27.0 mg/L^43^. Therefore, while our modeled DOC values are more accurate for values < 16 mg/L, predictions at rarely observed higher values have more error.

**Figure 7.**
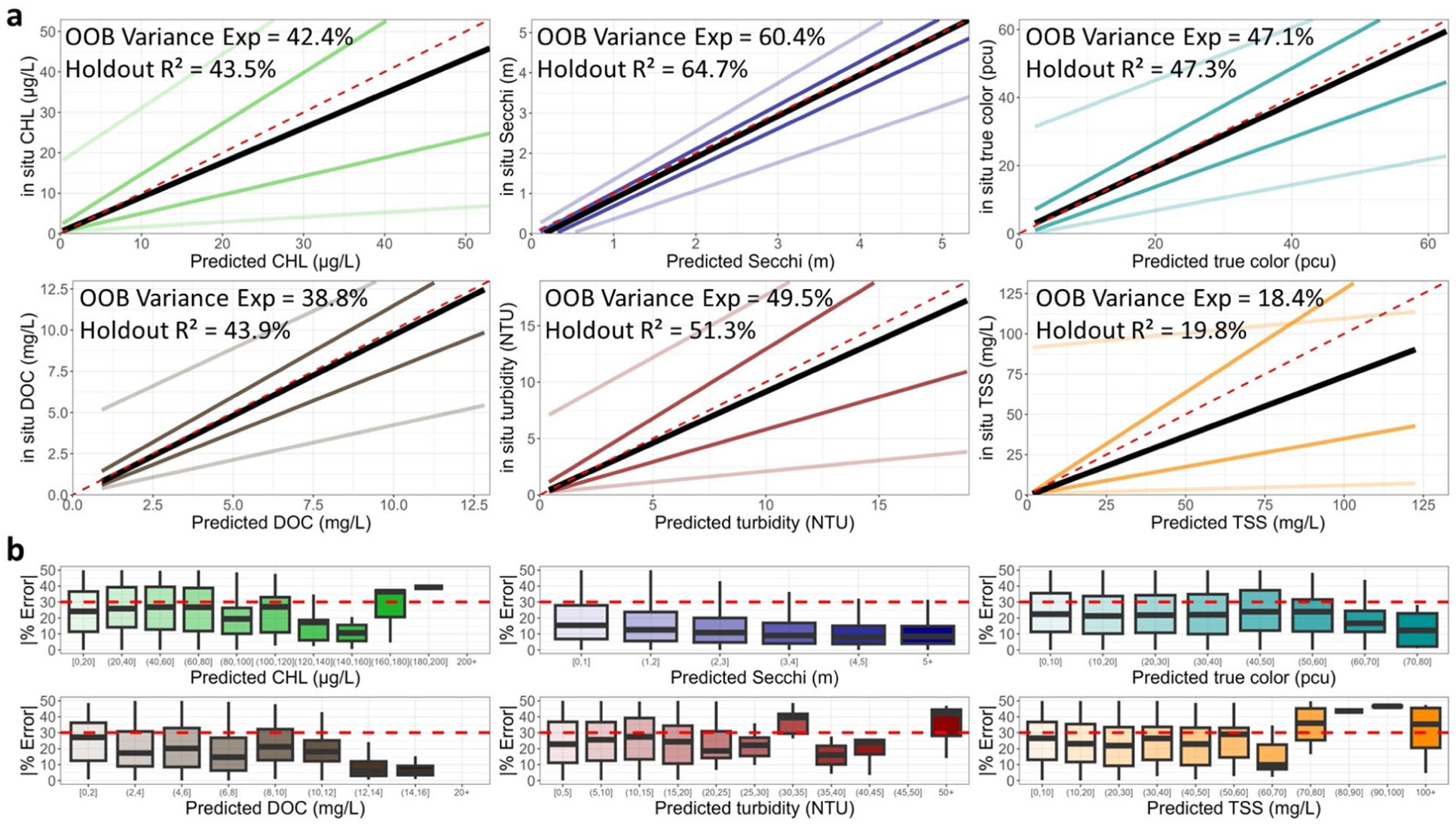
Fit statistic plots for the **Holdout-data model** of each of the six water quality variables in original units. The top six plots show predicted versus in situ quantile lines for the median quantile (black) and the 5th, 25th, 75th, and 95th quantiles (shaded colored lines). The maximum range of the X and Y axes is trimmed to the 90th percentile of in situ data (CHL = 53 ug/L; Secchi depth = 5.3 m; true color = 63 PCU; DOC = 13 mg/L; Turbidity = 19 NTU; TSS = 133 mg/L). The dashed red line is the 1:1 line. The internal random forest OOB variance explained from the model training data as well as the r-squared value of the holdout data are shown. The bottom 6 plots show the absolute percent error between predicted and in situ values in the model training dataset by intervals of predicted values for the training dataset. The red dashed line is the threshold identified as above which predictions are thought to be less reliable. Outliers are not plotted.

**Figure 8.**
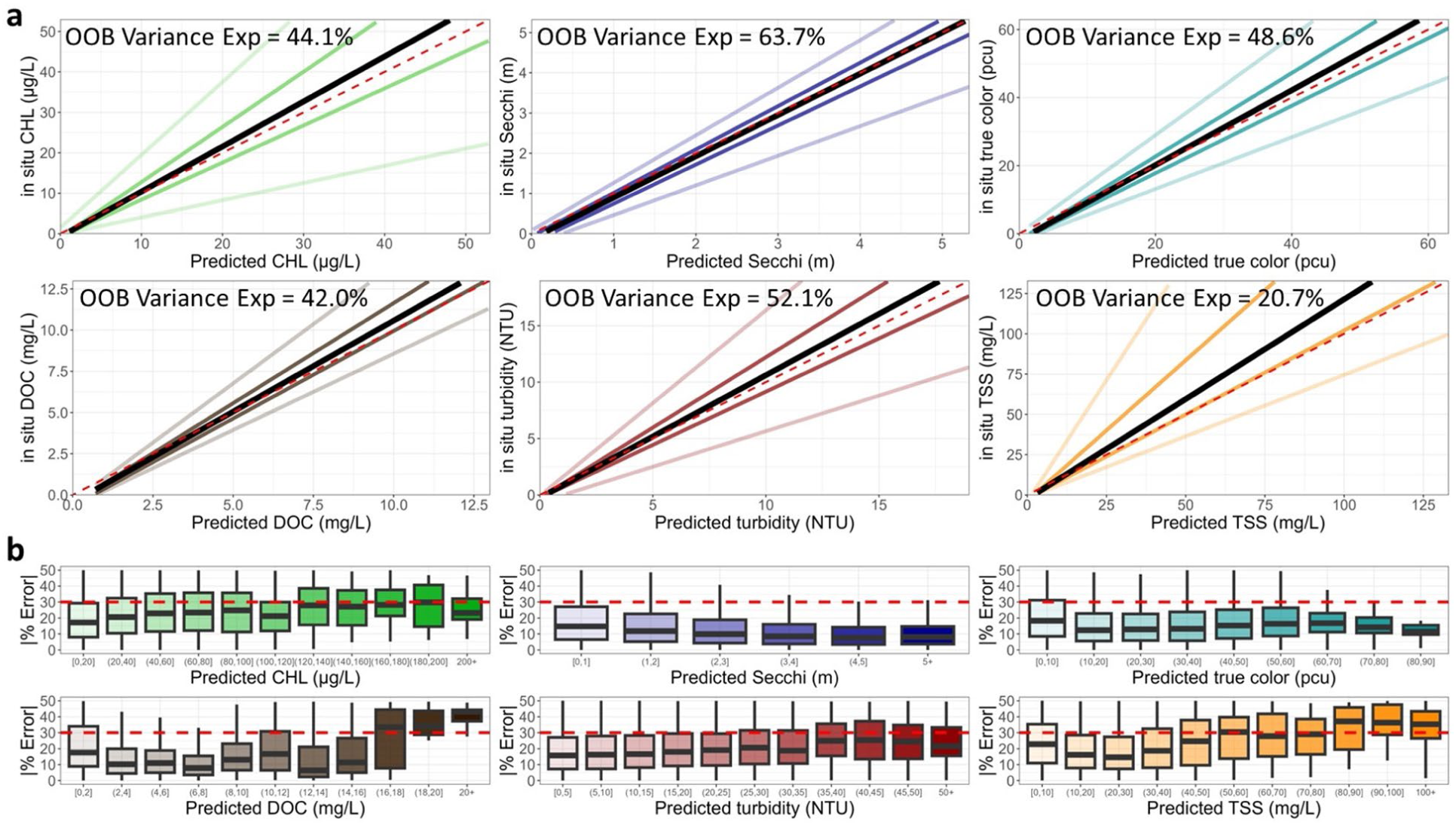
Fit statistic plots for the **Full-data predictive model** of each of the six water quality variables in original units. The top six plots show predicted versus in situ quantile lines for the median quantile (black) and the 5th, 25th, 75th, and 95th quantiles (shaded colored lines). The maximum range of the X and Y axes is trimmed to the 90th percentile of in situ data (CHL = 53 ug/L; Secchi depth = 5.3 m; true color = 63 PCU; DOC = 13 mg/L; Turbidity = 19 NTU; TSS = 133 mg/L). The dashed red line is the 1:1 line. The bottom 6 plots show the absolute percent error between predicted and in situ values in the model training dataset by intervals of predicted values for the training dataset. The red dashed line is the threshold identified as above which predictions may be less reliable.

Both sets of prediction errors provide similar conclusions where the OOB variance explained by the Holdout-data and Full-data models were substantially identical for each WQ variable. MAPE values were generally within the 30% range. Despite these differences, model predictions for the same 75% GRTS subset used in both frameworks were highly consistent, with R² values ≥ 0.92 across all WQ variables (**Figure 9b**). This consistency supports the reliability of both modeling approaches and affirms the utility of the dataset for large-scale ecological inference, provided users account for known sources of error using the supplied QA flags.

**Figure 9.**
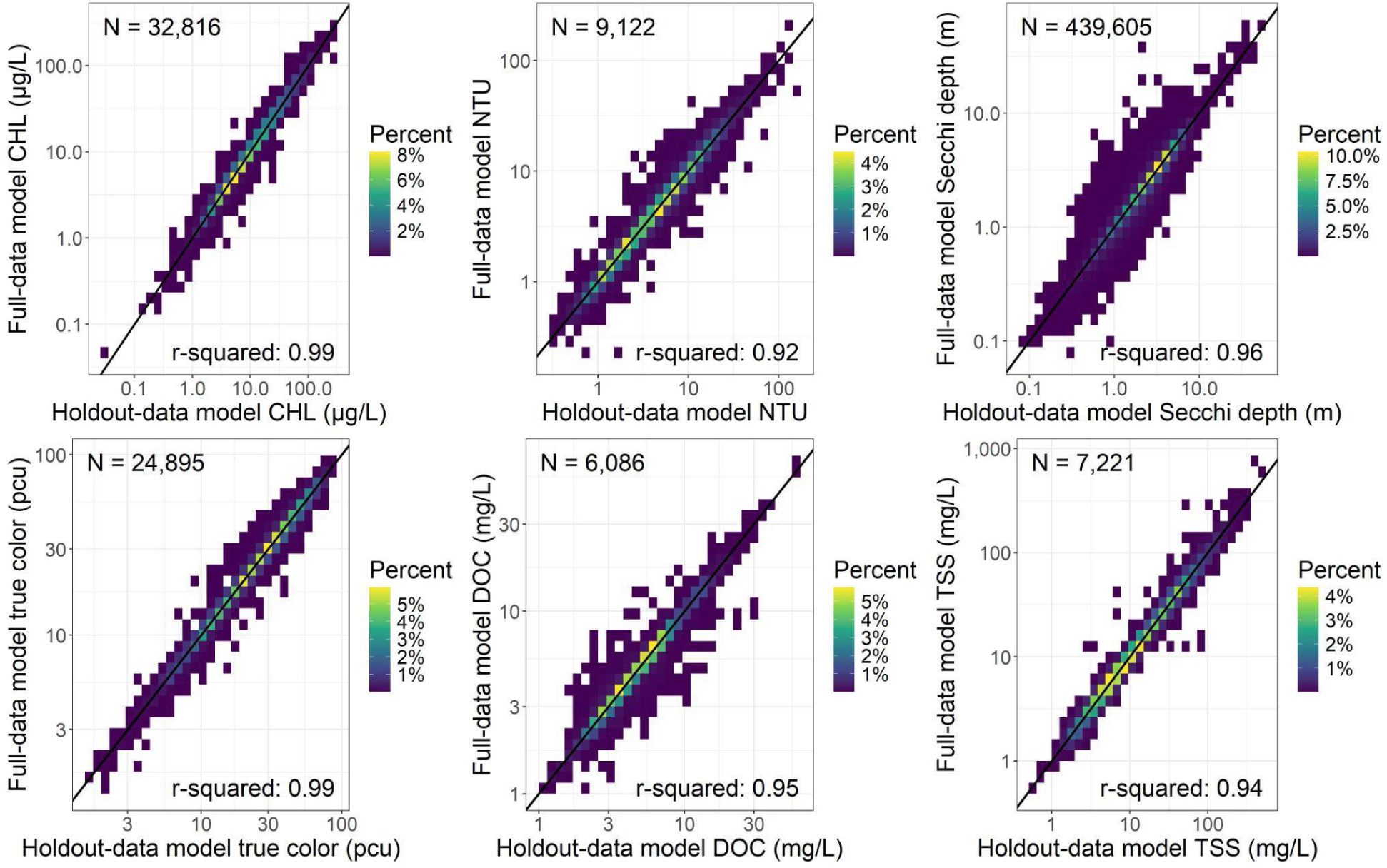
Comparison of the predictions from the Holdout-data and Full-data models for the same 75% of the data split into spatially representative test and train samples using Generalized Random Tessellation Stratified (GRTS) with the associated r-squared value for each water quality variable. The 1:1 line is represented by a solid black line.

### Model validation using individual lake data

To evaluate model performance at the individual lake scale, we analyzed lakes with dense, high-quality time series of in situ data and satellite imagery for three different WQ variables. First, we looked at Secchi depth data for Lake Okeechobee, a well-studied lake in Florida that contained 10,158 matchups within the ±7 window with satellite overpasses with 2,989 Secchi depth measurements. In this comparison, higher prediction errors occurred in concert with known issues with imagery data that we have flagged for in our database. Secchi depth was overestimated when the proportion of lake pixels available was low (< 10%; **Figure 10a**), an effect that is removed by eliminating such predictions from analysis. Second, in the same set of data from Lake Okeechobee, we found that negative reflectance values also led to prediction errors (**Figure 10 b-c).** Removing date-image combinations flagged with these errors results in the majority of the predicted versus in situ data falling close to the 1:1 line (**Figure 10d).**

**Figure 10.**
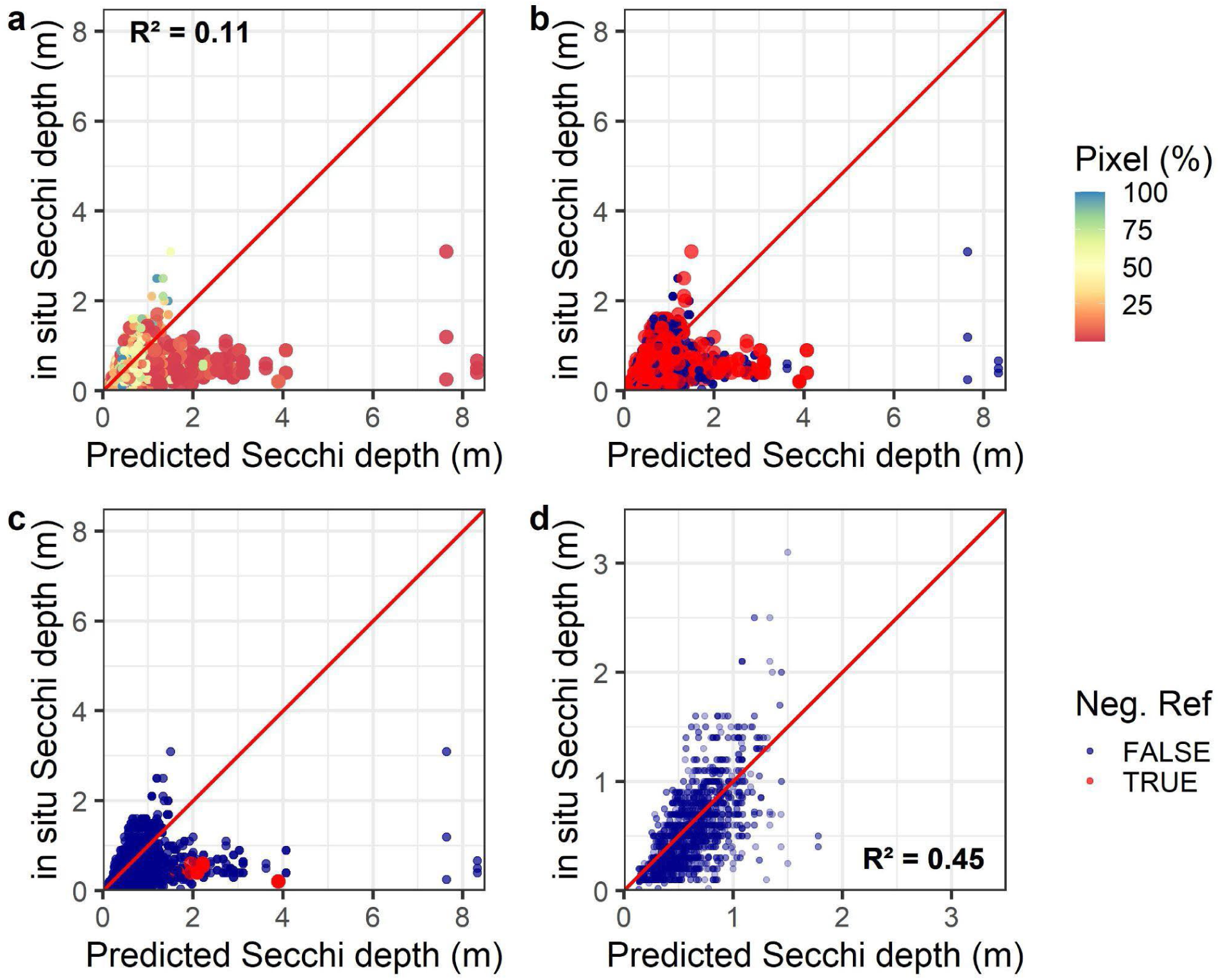
Analysis demonstrating pixel QAQC decisions and relationship to predictions. Predicted versus in situ Secchi depth (m) for a single lake in the dataset (Lake Okeechobee, FL) matchups in our training dataset (N = 2,989). (a) Predicted versus in situ Secchi depth (m) for all matchups color-coded by the percent of the lake’s pixels retrieved for that overpass out of the maximum ever retrieved across the 1984-2021 time series. (b) Predicted versus in situ Secchi depth coded by whether any satellite band for the overpass had a negative value for any lake pixel. (c) Predicted versus in situ Secchi depth coded by whether any satellite band for the overpass had a median reflectance value that was negative (note, this flag is part of the final prediction QAQC and is one of three components of our combined QAQC recommendation for good quality predictions). (d) Plot of predicted versus in situ Secchi depth after removing predictions with < 25% of a lake’s pixels retrieved or with negative median band reflectance. The 1:1 line is in red. R-squared values are presented prior to application of flags (a) and after removing predictions with < 25% of a lake’s pixels retrieved or with negative median band reflectance (d).

Further validation of the LAGOS-US LANDSAT dataset comes from a lake which has documented changes in water quality following a deliberate management intervention. The eutrophic Lake McCarrons in Minnesota was treated in 2004 with aluminum salts to reduce internal phosphorus loading, and, consequently, decreased algal growth^44^. The comparison between predicted and *in-situ* data for CHL suggested first, a close relationship near the 1:1 line (**Figure 11a**). Second, the alignment between the in situ data and predicted values improved when a narrower matchup window was applied (**Figure 11b-c**). However, the most striking was that the timing of the dramatic decrease in the in situ CHL was matched exactly in the predicted values from the remote-sensing models (**Figure 11d**). This lake thus presents an excellent case study to examine the application of our data product to individual lake studies beyond our primary goal of facilitating regional and continental-scale research.

**Figure 11.**
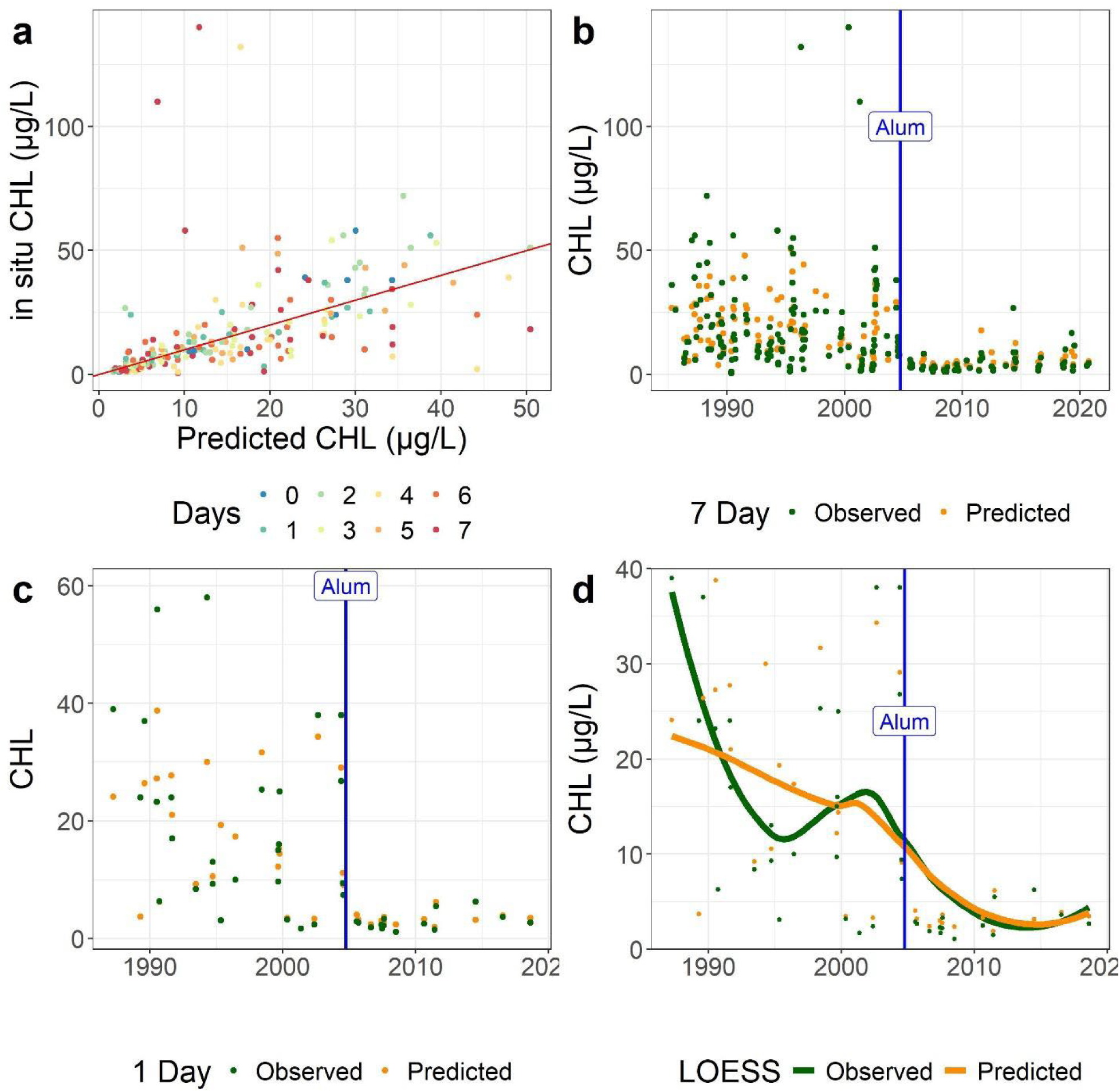
Predicted versus in situ CHL for Lake McCarrons, MN, which received alum treatment in October 2004. (a) Predicted versus in situ CHL across all matchup windows, color coded by the number of calendar days between the satellite overpass and the in situ measurement. The 1:1 line is in red. (b) The time series for the full matchup dataset for both in situ (green) and predicted (orange) CHL data. (c) The time series for the ±1 matchup dataset for both in situ (green) and predicted (orange) CHL. (d) Locally estimated scatterplot smoothing (LOESS) for the in situ and predicted CHL for the ±1 matchup dataset. The date of the alum treatment is shown as a vertical blue line for (c) and (d).

Finally, we demonstrate how our remote-sensing-based prediction dataset can be used to extend time series in lakes for which in situ data are missing. For example, DOC is not sampled as commonly as other water quality variables and yet is an important regulator of the ecology and biogeochemistry of inland waters, particularly related to carbon budgets^45^. Beaver Lake, Arkansas has a robust time series beginning in 2002 for in situ DOC data that aligns well with predicted DOC (**Figure 12a**). With our remote-sensing predictions, we can extend that time series back in time an additional 18 years (**Figure 12b**). These individual lake studies lend strong support for the potential for greatly expanding the estimates of water quality for individual inland lakes across the US in lakes with incomplete records or that have never been studied to date. Being able to estimate water quality for lakes across the entire US is an urgent priority to ensure that people have equitable access to good water quality regardless of socioeconomic status, race, ethnicity, and other factors. A recent study has shown that in situ estimates of water quality from government agencies and community science programs are not equitably distributed across all communities based on race and ethnicity^46^. Remotely sensed water quality predictions such as ours can help fill this gap in sampling.

**Figure 12.**
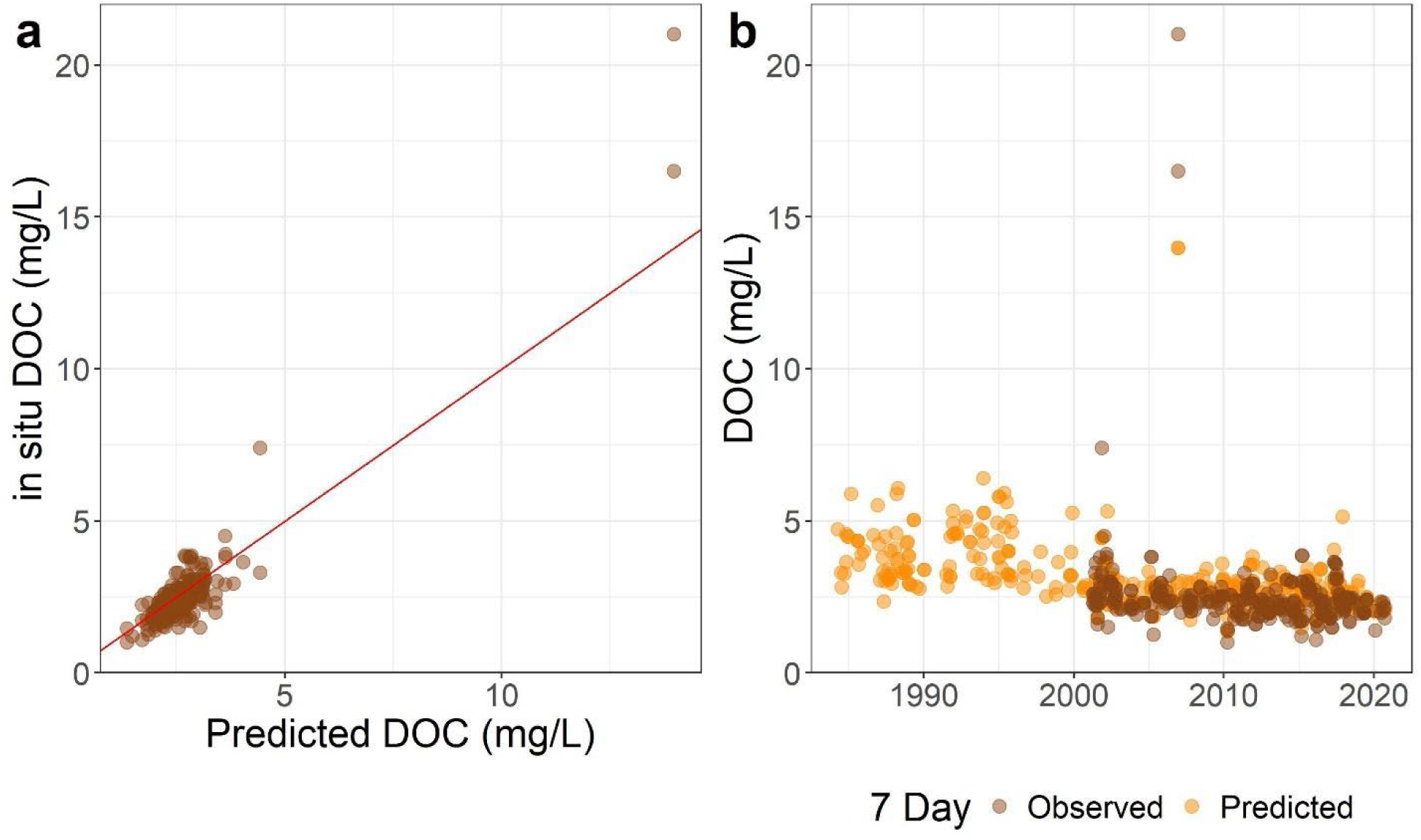
Plots showing DOC data for Beaver Lake, AR. (a) Predicted versus in situ DOC. (b) Predicted and in situ DOC through time.

## Usage Notes

### Data quality flags and filtering

In addition to the data values, our prediction dataset includes multiple measures of quality that provide users with the information needed to apply more lenient or stricter data quality filters: (1) if any pixel in any band for a lake retrieval had a negative reflectance value, (2) if any band for the whole lake retrieval had a negative median reflectance value, (3) the percent of pixels retrieved out of the maximum number of pixels ever retrieved for the lake across the prediction time series, and (4) if the retrieval for a lake shares the same calendar date as other retrievals due to overlapping scenes. The first two flags represent overcorrection over lake water pixels or other scene artifacts while the third provides an estimate of the extent of the lake surface represented in the predictions. Of all predictions, 3,145,912 (6.9%) were flagged for any negative reflectance pixel while 410,065 (0.9%) were flagged for median negative reflectance across the whole lake. Finally, because a lake can be within multiple Landsat row and path combinations due to scene overlap, we additionally flag predictions that have multiple retrievals for a single calendar date since users may want to treat these data differently than predictions for a single day. To summarize across these data flags, we include a variable called *QAQC_recommend* that identifies (with a value = TRUE) what we define to be research-ready and good quality data based on the following criteria: removal of retrievals with any median negative reflectance for any band, a threshold of least 10% of pixels being retrieved per lake, and removal of lower quality duplicate day predictions. The dataset filtered in this way includes a total sample size of 35,794,821 lake and date combinations and includes 78.0% of the total prediction sets. Another layer of quality control that users may want to consider is to drop predictions for DOC that exceed 16 mg/L and TSS that exceed 80 mg/L (**Figure 7**).

### Spatial and temporal distribution of predictions

The location and density of predictions for the six WQ variables are widely-distributed across the conterminous US (**Figure 13**). Due to cloudiness that is higher in the eastern US compared to the western US, the highest densities of prediction data occur in spatial banding patterns that result where Landsat overpasses overlap, and are more likely to contain cloud-free scenes that are known to occur with the Landsat platform^47^. A decrease in the spatial banding of the data can be seen from east to west and is likely a function of regional cloud patterns across the US. Because our dataset began with all lakes ≥ 4 ha, it is close to a census population of lakes, with the majority of lakes having at least 1 scene. Observations are also distributed across time (**Figure 14)**, with peak numbers of scenes occurring in April to May and then October through November - again, likely a result of general patterns of clouds. The number of images extracted through time increases through time as well, with the largest retrievals per year occurring after around 2015 that contain overlapping satellites that increase the potential for cloud-free scenes. Nevertheless, over 70,000 lakes across the US contain at least one summer water quality estimate across the full time period, resulting in an unprecedented time series dataset for lake WQ.

**Figure 13.**
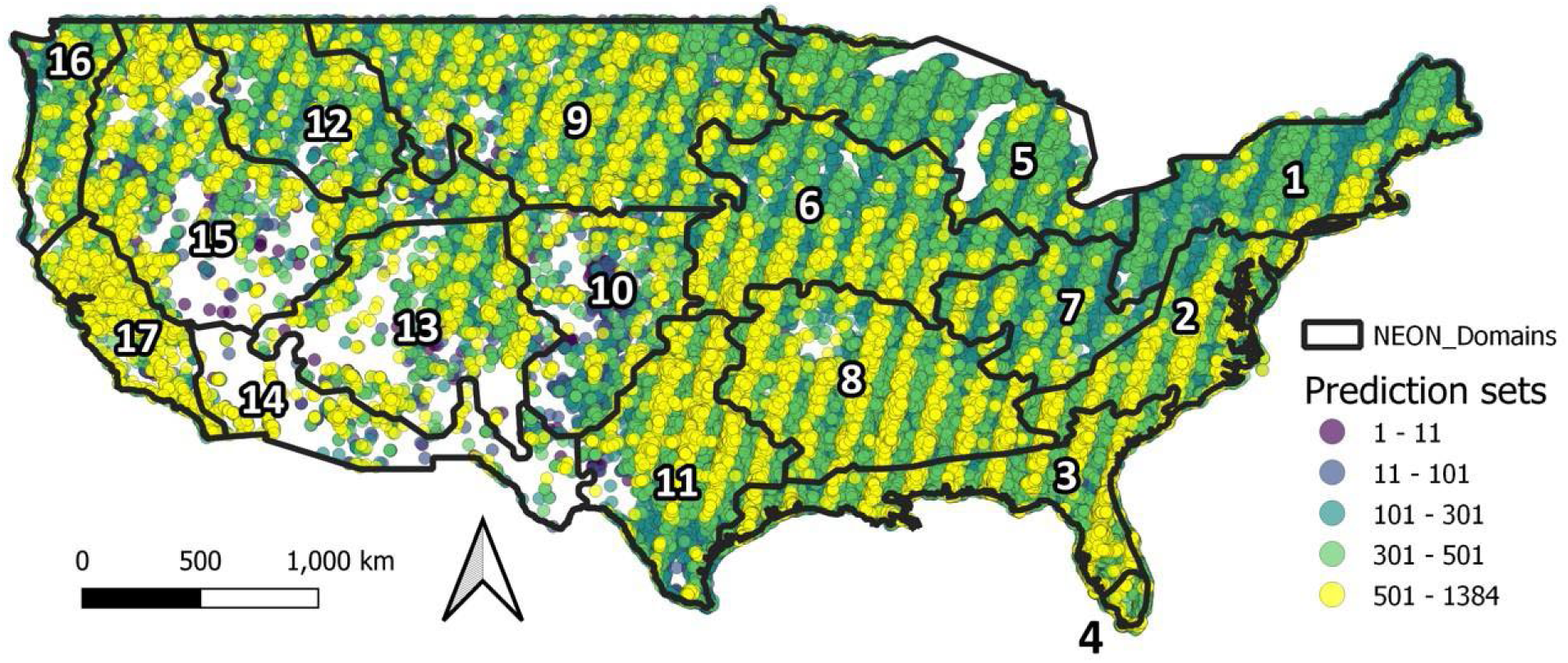
Map showing the total number of predictions for all 6 water quality variables by lake from 1984-2020. We include the prediction sets that have passed our QC filter (i.e., those without negative median reflectance for any band and that contain a minimum of 10% of the lake’s maximum pixels). The black borders show NEON region delineations.

**Figure 14.**
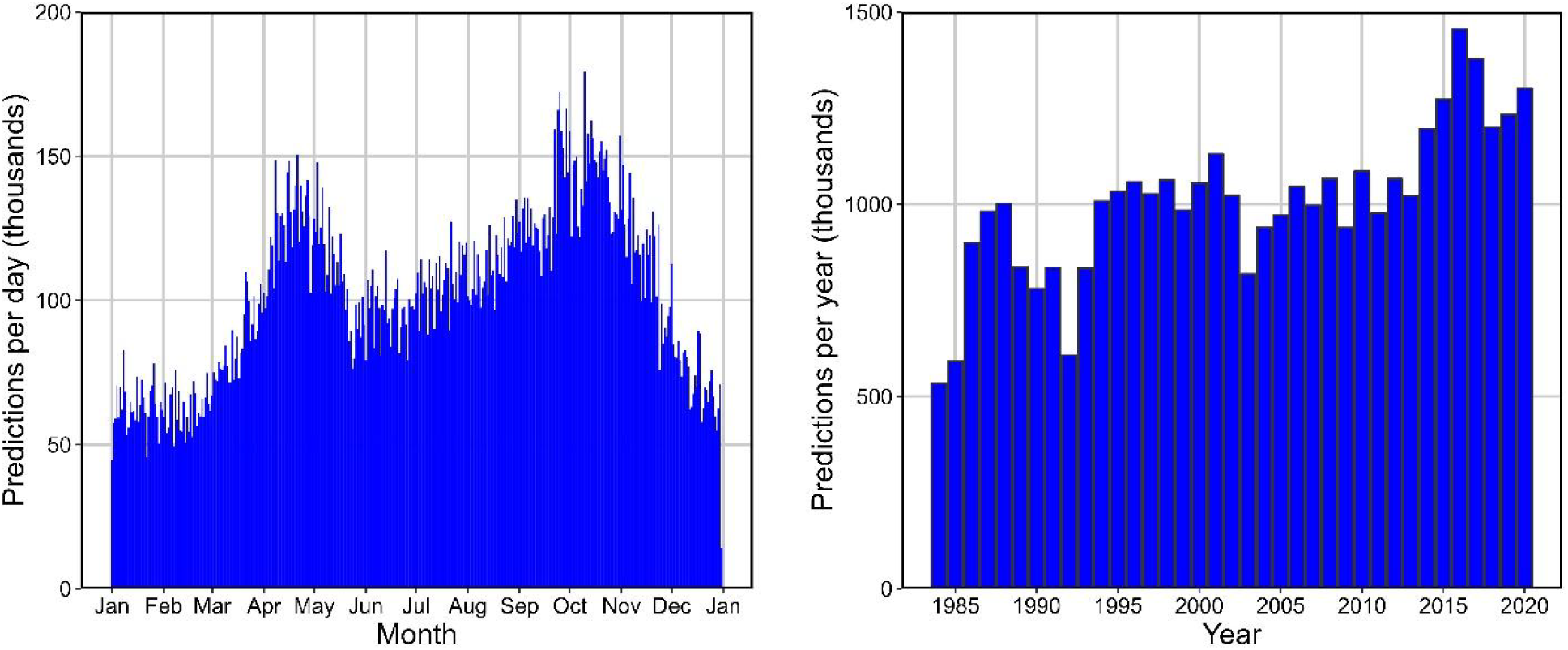
Plots showing the number of predictions in the final dataset, which applies to both predictive models individually. The left panel shows thousands of predictions per day by day of the year within each month. The panel on the right shows thousands of predictions by year. All data have had the minimum quality flags applied.

The ranges of the matchup dataset include broad ranges of all six WQ variables (**Figure 15**; **Table 3).** The prediction dataset generally had narrower distributions of the data as shown by the quantile plots that showed less spread in the data. However, given that we know that most lakes in the US have never been sampled (Shuvo et al. 2023), we would not have expected the ranges of the matchup dataset to match the prediction dataset closely given the biases in water quality sampling programs^48,49^. Examining sampling sizes through time is also valuable. The median number of observations per lake is 257 (**Table 3**); the median lake-year combination with WQ observations is 37; and the median number of lakes per year with observations is 121,812.

**Figure 15.**
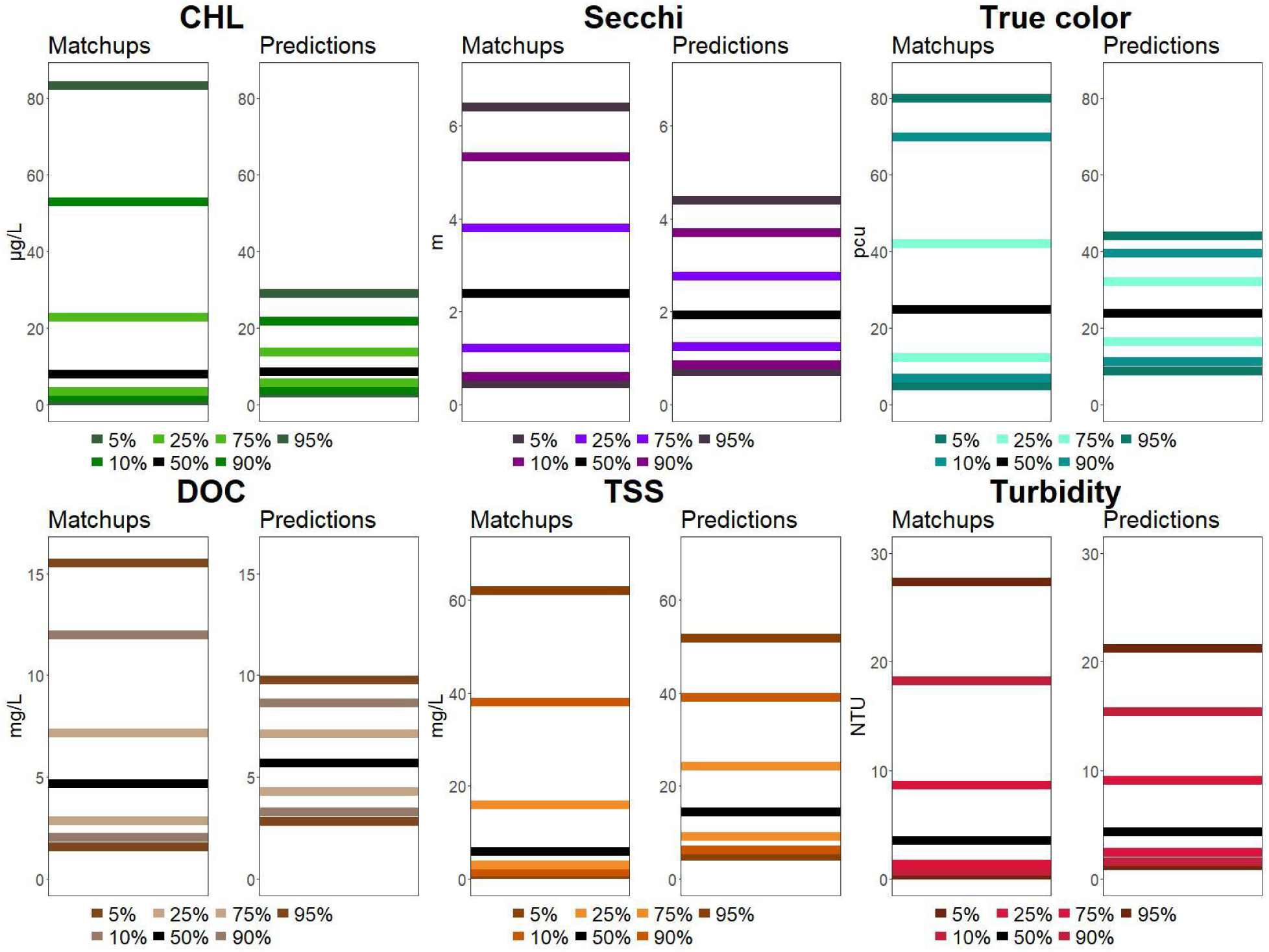
Plots of quantiles for each of the six water quality variables from the matchup dataset (left plots) and from the Full-data model prediction dataset (right plots). Black lines are median values.

**Table 3.** Statistical summary table of predictions for the six water quality variables, along with summaries of predictions per lake, and per year. Lake area was from LAGOS-US LOCUS^23^.

| Variable | Units | min | q5 | q25 | q50 | q75 | q95 | max |
| --- | --- | --- | --- | --- | --- | --- | --- | --- |
| *Water quality variables (37,251,456 predictions)* |  |  |  |  |  |  |  |  |
| **CHL** | µg/L | 0.04 | 3.1 | 5.7 | 8.7 | 14.0 | 29.1 | 299.9 |
| **True color** | PCU | 0.1 | 8.7 | 16.4 | 23.9 | 32.4 | 44.7 | 88.1 |
| **DOC** | mg/L | 0.5 | 2.8 | 4.3 | 5.7 | 7.1 | 9.7 | 67.7 |
| **Turbidity** | NTU | 0.05 | 1.2 | 2.4 | 4.4 | 9.3 | 21.2 | 197.8 |
| **Secchi depth** | meters | 0.02 | 0.7 | 1.2 | 1.9 | 2.7 | 4.3 | 59.3 |
| **TSS** | mg/L | 0.5 | 4.8 | 9.1 | 14.2 | 24.3 | 51.6 | 675.9 |
| *Lake-related (136,877 lakes)* |  |  |  |  |  |  |  |  |
| **Predictions per lake** | N | 1 | 32 | 160 | 255 | 361 | 568 | 1377 |
| **Lake-years with predictions\*** | N | 1 | 13 | 32 | 37 | 37 | 37 | 37 |
| **Lake water area** | ha | 4 | 4 | 6 | 9 | 19 | 116 | 338,950 |
| *Year-related (37 years)* |  |  |  |  |  |  |  |  |
| **Number of lakes per year** | N | 108,158 | 110,454 | 117,689 | 121,821 | 123,719 | 125,928 | 126,177 |
| **Number of predictions per year** | N | 534,405 | 604,508 | 938,970 | 1,021,304 | 1,066,120 | 1,317,022 | 1,455,641 |

\*The percentages of lakes with 37 and 30+ years with at least 1 summer observation (June-September) are, respectively, 14.8% and 51.6%

### Water quality variable distributions and correlations

Although there is overlap in reflectances that these six WQ variables are related to (e.g, overlapping absorption spectra for CDOM and chlorophyll documented by Olmanson et al. 2015), the dominant factors that these WQ variables represent are distinct enough that they are predicted by different bands and band ratios (**Figure 6**), and, despite very high sample sizes, they are not highly correlated with the exception of Secchi depth which is highly correlated to everything except true color (**Figure 16**). In fact, these correlations match expected ecological relationships. For example, Secchi depth and turbidity, both measures of water clarity, are the most strongly correlated in the dataset (r=0.90). In contrast, the correlation between true color, which measures dissolved fractions, and TSS, which measures suspended particles, is the lowest among all comparisons (r=0.3).

**Figure 16.**
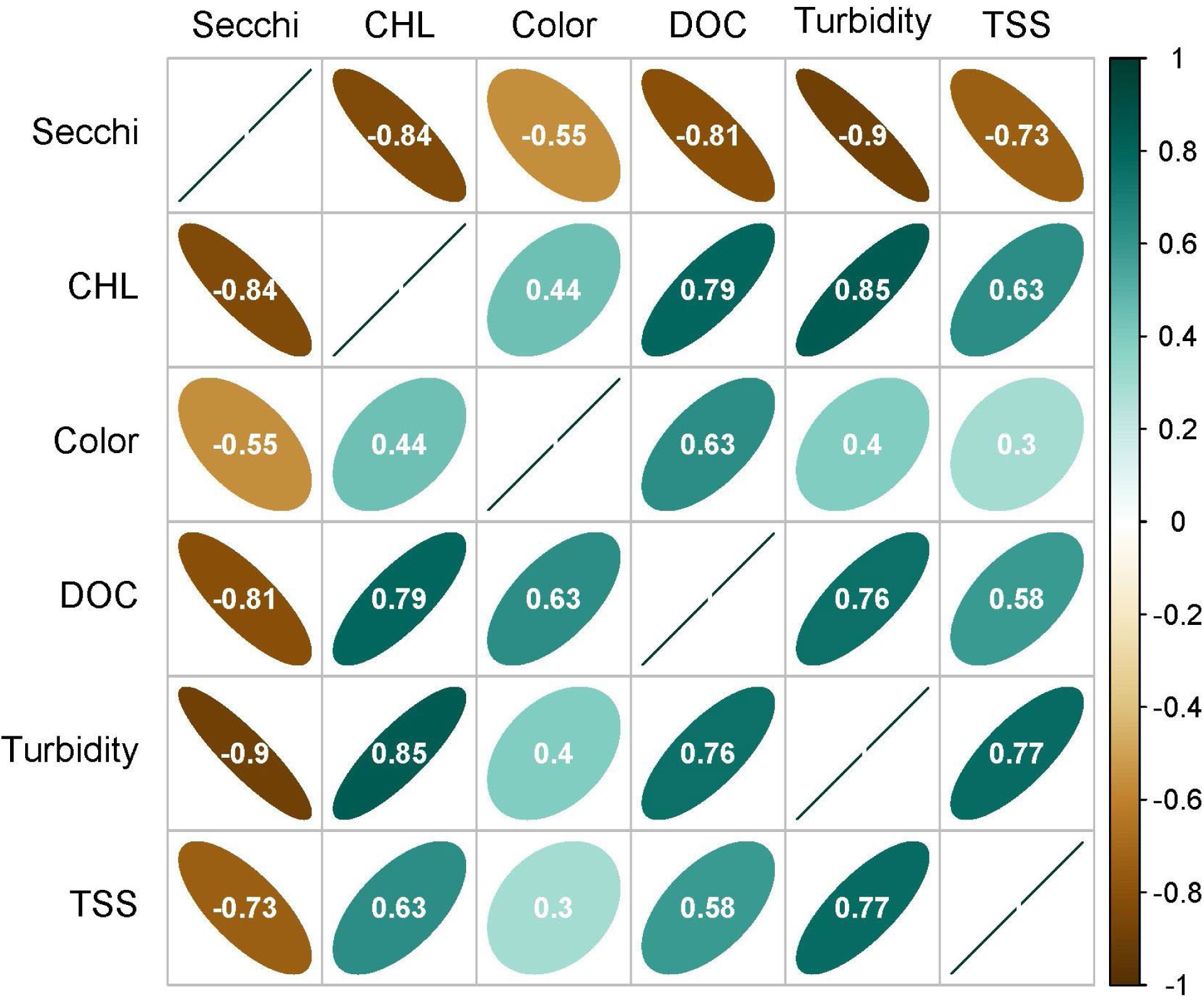
Correlation matrix of predicted values from the Full-data model for the six water quality variables. Data points were randomly selected single summer (June - September) predicted values for each year from 1984-2020 for the 20,191 lakes with 37 years of data. Note, all WQ variables were from the same scene for a lake year. Spearman rank correlations were based on log_10_ transformed data. All p values << 0.00001. Color is true color.

To examine relationships among the variables further, we also applied a common threshold for eutrophication that separates oligo-mesotrophic lakes from eu-hypereutrophic lakes (CHL ≤ 7 ug/l) and examined relationships among CHL, Secchi depth, and DOC. The patterns (**Figure 17)** make ecological sense based on our current understanding of key limnological relationships ^28,50^. First, DOC is positively related to true color in both sets of lakes, but DOC at a given color value tends to be lower in oligo-mesotrophic lakes than in eu-hypereutrophic lakes. These latter lakes have the highest DOC values in the dataset, particularly where with true color > 20 suggesting high contributions from both non-colored DOC from high algal growth and colored DOC from tannins that occur in high color waters. Second, Secchi depth versus true color also diverges in the lakes in the two trophic classes, with a far clearer relationship between the two variables in oligo-mesotrophic lakes compared to eu-hypereutrophic lakes suggesting a shift in control of water clarity from color to algal biomass. Finally, the importance of Secchi depth and its relationships with all other variables is demonstrated in **Figure 18**. Secchi depth is the most commonly sampled WQ variable in US lakes^10^, and its inclusion in nutrient modeling of lakes has been found to decrease prediction error^51,52^. Being able to predict Secchi depth in an even broader range of lakes has great potential for improving water quality modeling at broad scales.

**Figure 17.**
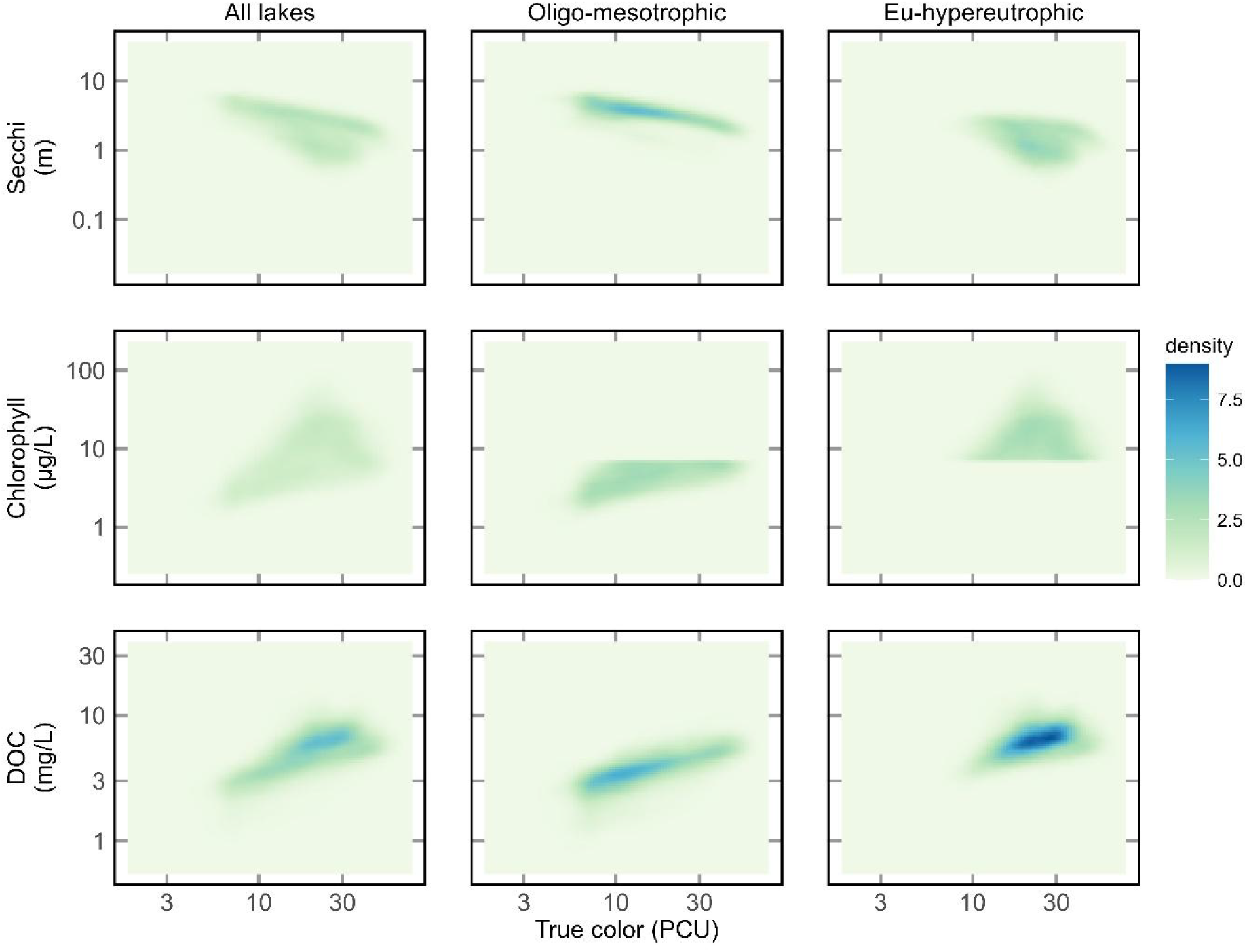
Density plots of predicted values from the Full-data model for true color plotted against three of the water quality variables related to nutrient and color status. Both axes are plotted on a log10 scale. Data points were randomly selected single summer (June - September) predictions for each year from 1984-2020 for the 20,191 lakes with 37 years of data. Note, all WQ variables were from the same scene for a lake year. Columns represent all lakes and lakes were placed into two trophic classifications based on CHL with values ≤ 7 ug/L classified as oligo-mesotrophic and values > 7 ug/L as eutrophic-hypereutrophic.

**Figure 18.**
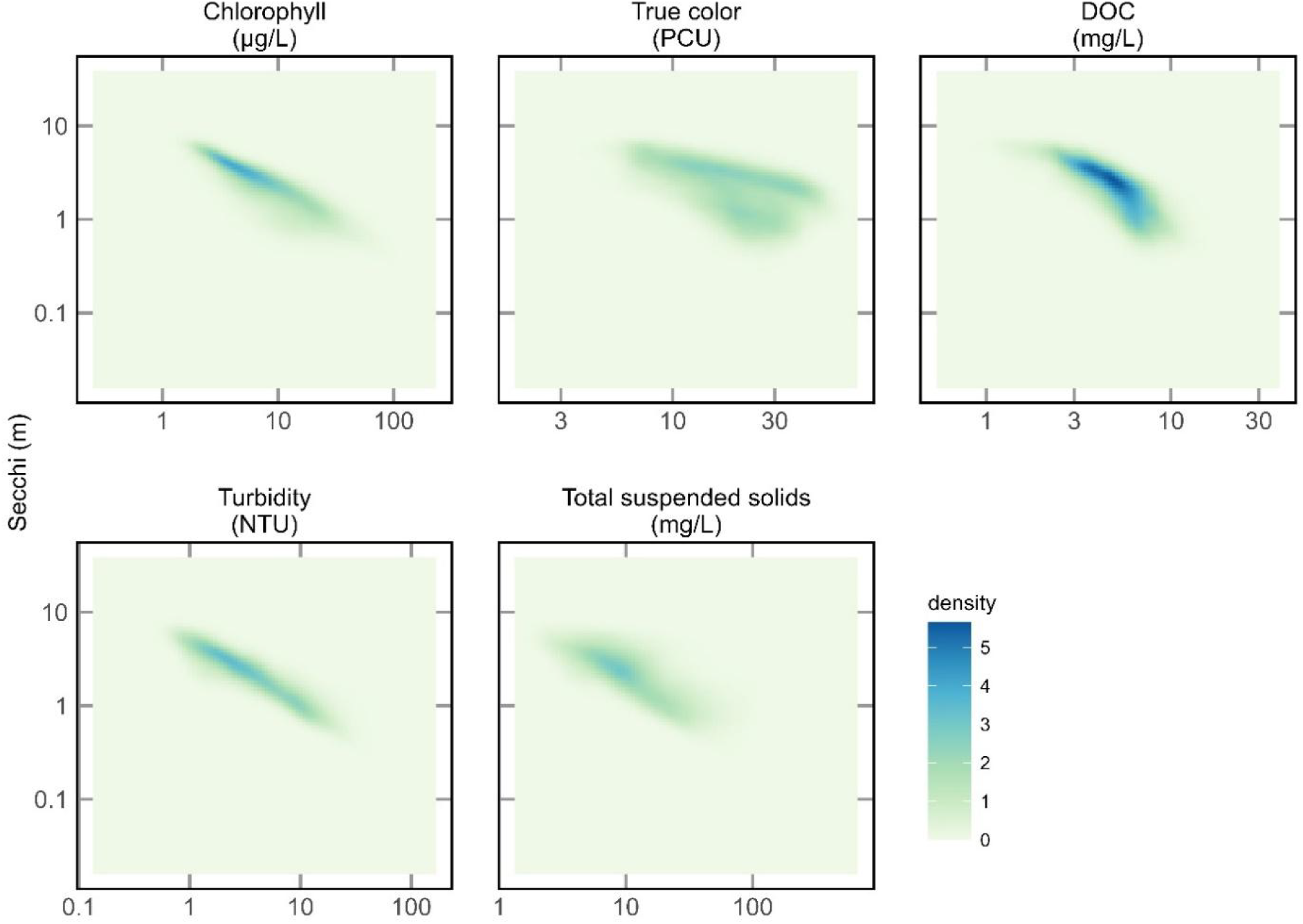
Density plots of Secchi depth against all other water quality variables from the Full-data model predictions. Both axes are plotted on a log10 scale. Data points were randomly selected single summer (June - September) predictions for each year from 1984-2020 for the 20,191 lakes with 37 years of data. Note, all WQ variables were from the same scene for a lake year.

### Filtering and combining predictions for temporal analysis

Users can filter and combine LAGOS-US LANDSAT predictions to support diverse modeling frameworks and research questions. For example, in Soranno et al.^53^, retained only lake-date combinations with the highest quality data based on the QAQC_recommend flag, and lakes that had at least eight valid chlorophyll-a observations per summer over a 34-year period, excluding lakes with more than one consecutive year missing. These criteria resulted in annual summaries of lake CHL in 24,452 lakes that were used in advanced time series models that require dense, temporally regular observations. Therefore, users conducting time series analyses to address other types of research questions might consider: (1) setting minimum observation thresholds per lake-year, (2) excluding pre-2000 data due to lower scene availability, (3) restricting analyses to ice-free months (e.g., April–October), and (4) applying the QAQC_recommend flag for highest quality predictions. The data density of predictions per year for lakes is shown in **Figure 19**.

**Figure 19.**
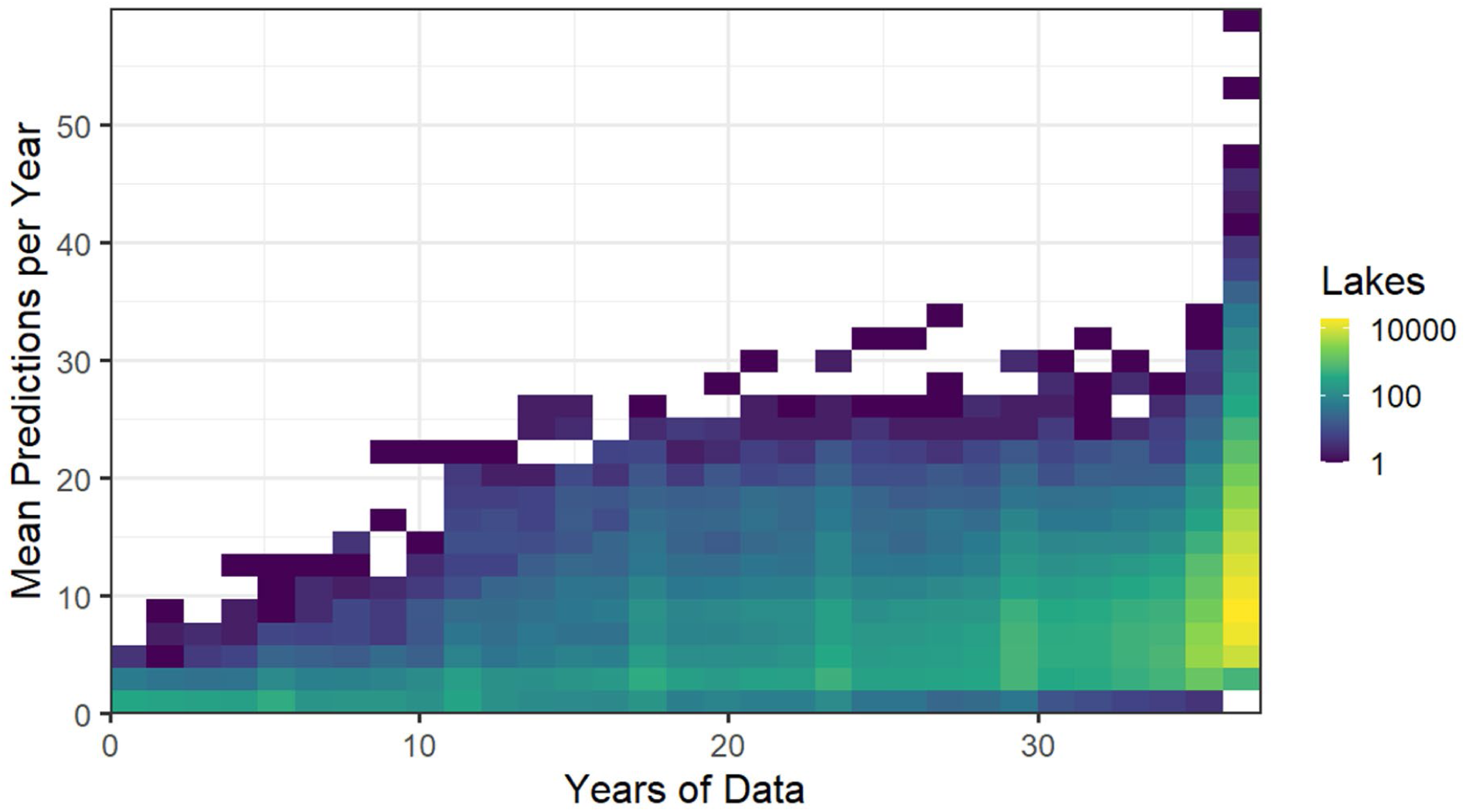
Density plot of the number of lakes for a given number of mean predictions per year and the total number of years of data. These values represent the maximal data amount for lakes in the prediction datasets prior to the application of QAQC flags. The Full-data and Holdout-data models have the identical number of predictions presented here.

### Summary

LAGOS-US LANDSAT is the first published dataset of remotely-sensed, ground-truthed estimates of multiple WQ variables providing near-census coverage of lakes ≥4 ha (>100,000 lakes) at the continental scale that also includes:

● Coverage of small lakes (≥4 ha), with 50% of observations from lakes <9 ha
● Whole-lake water pixel extractions of all bands
● Complete workflow products in a single, integrated dataset including data quality flags
● Full interoperability with the LAGOS-US research platform

The LAGOS-US platform provides comprehensive geospatial and temporal datasets for macroscale lake studies (watershed delineations, land use/cover, geology, soils, climate, stream connections). This LANDSAT module adds 37 years of water quality data on > 100,000 lakes, enabling unprecedented long-term analyses of lake ecosystems at macroscales.

## Code Availability

The entire code, including the code used for extracting reflectances in Google Earth Engine, is available in Zenodo at https://doi.org/10.5281/zenodo.1111028354

## Acknowledgments

We thank the many contributors who created the LAGOS-US research platform, especially those researchers who created the LAGOS-US LIMNO data module, and the original data providers for this rich source of in situ data without which this data product would not be possible. This research was funded by the National Science Foundation (DEB-1638679). PAS also acknowledges support from the National Institute of Food and Agriculture, Hatch Project 176820. This material is based upon work supported by (while serving at) the National Science Foundation from 2019-2023 for PAS on her independent research and development time. Thanks also to Yvonne Allen, Spatial Ecologist, US FWS-Southeast Region for the idea of using the Google Earth Engine platform for inland aquatic studies of water quality parameters and for a very helpful review of an earlier draft.

## Author contributions

PJH had the initial idea and developed and implemented the analytical approach for this dataset. All authors contributed to making decisions about the modeling approach as well as outlining and writing the article. PJH created all figures and tables except for Figure 1 and Table 1-2 (PAS) and Figure 14, 16-18, and Table 3 (KEW).

